# Open and closed forms of assembled henipavirus nucleoprotein suggest structural basis of genome access

**DOI:** 10.1101/2025.11.02.686081

**Authors:** Rupesh Balaji Jayachandran, Erwan Quignon, Max Renner

## Abstract

Henipaviruses, such as Nipah virus, can cause deadly illness and constitute WHO blueprint priorities due to their pandemic potential. Their genomes are packaged within a nucleocapsid consisting of viral nucleoproteins (N). Currently, it is unclear how the encapsidated genome is released from N to allow the viral polymerase to read its sequence. Here, we present the first high-resolution cryo-EM structure of a helical N-RNA filament from Langya henipavirus (LayV), allowing us to identify vertical interactions crucial for assembly. We show that assembly efficiency is sequence-dependent and prefers 5’-genomic sequences. Further, we solve the structure of an RNA-free assembly of LayV N. Structural comparison of the RNA-bound and RNA-free LayV N shows a conformational opening and closing, even within the assembled state. Our data suggest that N within nucleocapsids may undergo local conformational changes, switching between closed and open states, to temporarily allow access to the encapsidated RNA without nucleocapsid disruption.

## Introduction

Langya henipavirus (LayV) is an emerging henipavirus that was identified in 2022 in eastern China where it was isolated from patients with febrile illness (*1*). LayV is closely related to Nipah virus (NiV) and Hendra virus (HeV), the two prototypic henipaviruses (*2*). Since their discovery in the 1990s (*3*, *4*), NiV and HeV have been responsible for regional outbreaks in parts of South and Southeast Asia and in Australia, respectively (*5*). NiV outbreaks are often deadly with case fatality rates of >60% (*6*), and involving human-to-human transmission events (*7*, *8*). Symptoms of henipaviral disease include fever, myalgia, cough, and in fatal cases, severe respiratory illness and encephalitis (*6*).

LayV is the latest henipavirus to be associated with human infections after NiV, HeV, and the Mòjiāng virus (MojV), the latter of which may be associated with fatal cases in China (*9*). Unlike the bat-borne NiV and HeV (*10*, *11*), LayV and MojV are thought to have spilled over from shrews and rats, respectively (*1*, *9*). In recent years, further rodent-borne henipaviruses were isolated in China (*12*, *13*), South Korea (*14*), Belgium (*15*, *16*), and Guinea (*15*), which led to the creation of a new genus called *parahenipavirus* to distinguish the rodent-borne henipaviruses from those transmitted by bats (*17*) (Supplementary Figure 1A). The lack of approved vaccines or treatment options available for human use against henipaviruses, the expanding geographical range of animal hosts, and diverse species tropism exhibited by henipaviruses have led the WHO to include henipaviral diseases in its blueprint list of priority diseases (*18*).

Like other members of *Paramyxoviridae* (see phylogenetic tree in Supplementary Figure 1B), LayV is a non-segmented, negative-strand RNA virus (nsNSV) belonging to the order of *Mononegavirales*. The (-ss)RNA genome of LayV is ~18 kb long and it contains six genes, which encode the nucleoprotein (N), phosphoprotein (P), matrix protein (M), fusion protein (F), glycoprotein (G), and large protein (L) (*1*). Additionally, the open reading frame of P also encodes the non-structural proteins V, W, and C (*1*). Cell attachment and entry are mediated by the two surface glycoproteins G and F respectively (*19*, *20*) while M is crucial for the assembly and budding of new virions (*21*). Transcription and replication of the RNA genome are facilitated by the RNA synthesis machinery of the virus, comprising the polymerase complex and the nucleocapsid. The polymerase complex consists of the multifunctional L and its obligate cofactor, P, which forms a homotetramer, P_4_ (*22*). N encapsidates the RNA genome (and antigenome) to form the nucleocapsid which acts as the template for both transcription and replication. The nucleocapsid protects the RNA genome from host nucleases and prevents innate immune signalling (*23*).

*Mononegavirales* transcription and replication take place in membrane-less organelles called ‘inclusion bodies’, which are formed within the cytoplasm of the host cell (*24*, *25*). Within inclusion bodies, the polymerase complex scans the encapsidated RNA template for *cis*-acting elements and initiates *de novo* RNA synthesis (*26*). Initially, the polymerase complex mediates transcription to produce capped and polyadenylated viral mRNAs in a descending gradient from 3’ end to 5’ end, with N-encoding mRNAs being the most abundant. Viral mRNAs are then translated by the host-cell machinery. Once the concentration of N reaches a threshold level, the polymerase complex switches to genome replication, where it produces complementary antigenomes acting as templates for synthesis of further genomes (*27*). During genome synthesis, the nascent RNA is concomitantly encapsidated by monomeric N, forming a progeny nucleocapsid. Paramyxoviruses possess helical nucleocapsids of left-handed chirality that bear a characteristic ‘herringbone’ appearance (*28–31*).

The formation of helical, nucleocapsid-like assemblies can occur with recombinant paramyxoviral N (*32–36*). In addition, depending on virus and expression system, ring- (*35*, *37*) and clam- (*36*, *38–40*) shaped oligomers have been observed. Irrespective of the structural arrangement of N-assemblies, N binds exactly six nucleotides of RNA. This is the basis of the so-called ‘rule of six’, a feature unique to paramyxoviruses, requiring the number of bases in their genomes be multiples of six (*32*, *41*). The six nucleotides per protomer are sequestered in a positively charged binding site, located at the interface formed by the two globular domains of the ordered core of N (N_core_) – the N-terminal (N_NTD_) and C-terminal domains (N_CTD_). In presence of RNA, the N_NTD_ and N_CTD_ change conformation, tilting towards each other and forming a tight RNA-binding cleft. The N_core_ in its RNA-bound form with the N_NTD_ and N_CTD_ clamping onto the RNA is thus termed to be in a ‘closed’ conformation. In the absence of RNA, the N_NTD_ and N_CTD_ are shifted away from each other, in what is known as the ‘open’ conformation of N_core_ (*23*). In addition to N_core_, N also possesses an extended disordered tail (N_tail_), which mediates interactions with the polymerase complex through P (*42*), and NT- and CT-arms that extend from the N_NTD_ and N_CTD_, respectively.

Monomeric N is unstable on its own, unless bound to the N-terminal domain of P (P_NTD_) which acts as a chaperone for monomeric and RNA-free N in complexes known as N^0^-P (*23*). Among paramyxoviruses, N^0^-P structures have been characterized in NiV, measles virus (MeV), parainfluenza virus 5 (PIV5), and human parainfluenza virus 3 (hPIV3) (*43–46*). Structural comparisons of RNA-bound N assemblies and RNA-free N^0^-P complexes for MeV, PIV5, and NiV have helped shape our understanding of how N monomers assemble to nucleocapsids during replication by L. Despite these insights, fundamental mechanistic questions concerning transcription and replication remain. A widespread model, the so-called ‘cartwheeling model’, has been proposed to explain how the polymerase complex moves along the nucleocapsid (*47*). In this model, the oligomeric P_4_ cartwheels along the template while transiently binding and unbinding to the N_tail_ of adjacent N protomers, an interaction mediated by the X-domain of P (P_XD_) (*48*, *49*). While the model explains how the polymerase complex ma>y traverse along the length of the nucleocapsid, it remains wholly unclear how the occluded bases of the nucleocapsid are extracted such that they can serve as template for RNA synthesis.

Here, we present the first high-resolution cryo-EM structure of a helical N-RNA assembly from a henipavirus, LayV-N, at 3.1 Å resolution. We assemble the structure *in vitro* from purified N monomers by incubation with RNA. Using time-resolved fluorescence anisotropy, we observe that the assembly efficiency of helical N-RNA from monomers is sequence dependent and prefers 5’ genomic sequences. While lateral contacts have been described previously for N oligomers, we are now able to characterize vertical interactions, crucial to form the helical structure. Further, we solve the cryo-EM structure of a ring-like assembly of LayV-N at 2.6 Å resolution. Surprisingly, and in contrast to previous studies, the 13-mer N assembly is absent of RNA. Structural comparison of the RNA-bound and RNA-free LayV-N shows a conformational opening and closing of N_core_, even in the absence of P and in the assembled state. Our data suggest that N within paramyxoviral nucleocapsids may undergo local conformational changes, switching between closed and open states, to temporarily allow access to the encapsidated RNA without disrupting the nucleocapsid.

## Results

### LayV-N_core_ exists in an equilibrium between its oligomeric and monomeric states in solution

Paramyxoviral N, produced in bacterial expression systems, has been widely used for structural studies (*33–40*). The resulting assemblies typically possess well-resolved N_core_ while bound to host RNA and no observable density for N_tail_, even when the full-length protein is expressed (*35–40*). The lack of resolved density for N_tail_ is indicative of its flexible nature. Removal of N_tail_ resulted in the formation of more rigid helical packing in SeV, compared to full-length N (*39*). To enhance the possibility of acquiring well-resolved N assemblies, we designed a LayV-N expression construct without the N_tail_, referred to as LayV-N_core_ hereafter (Figure 1A). The N_tail_ (residues 421-539) region was identified based on a disorder score plot of full-length LayV-N (Supplementary Figure 2A) using Metapredict v3 (*50*).

**Figure 1:**
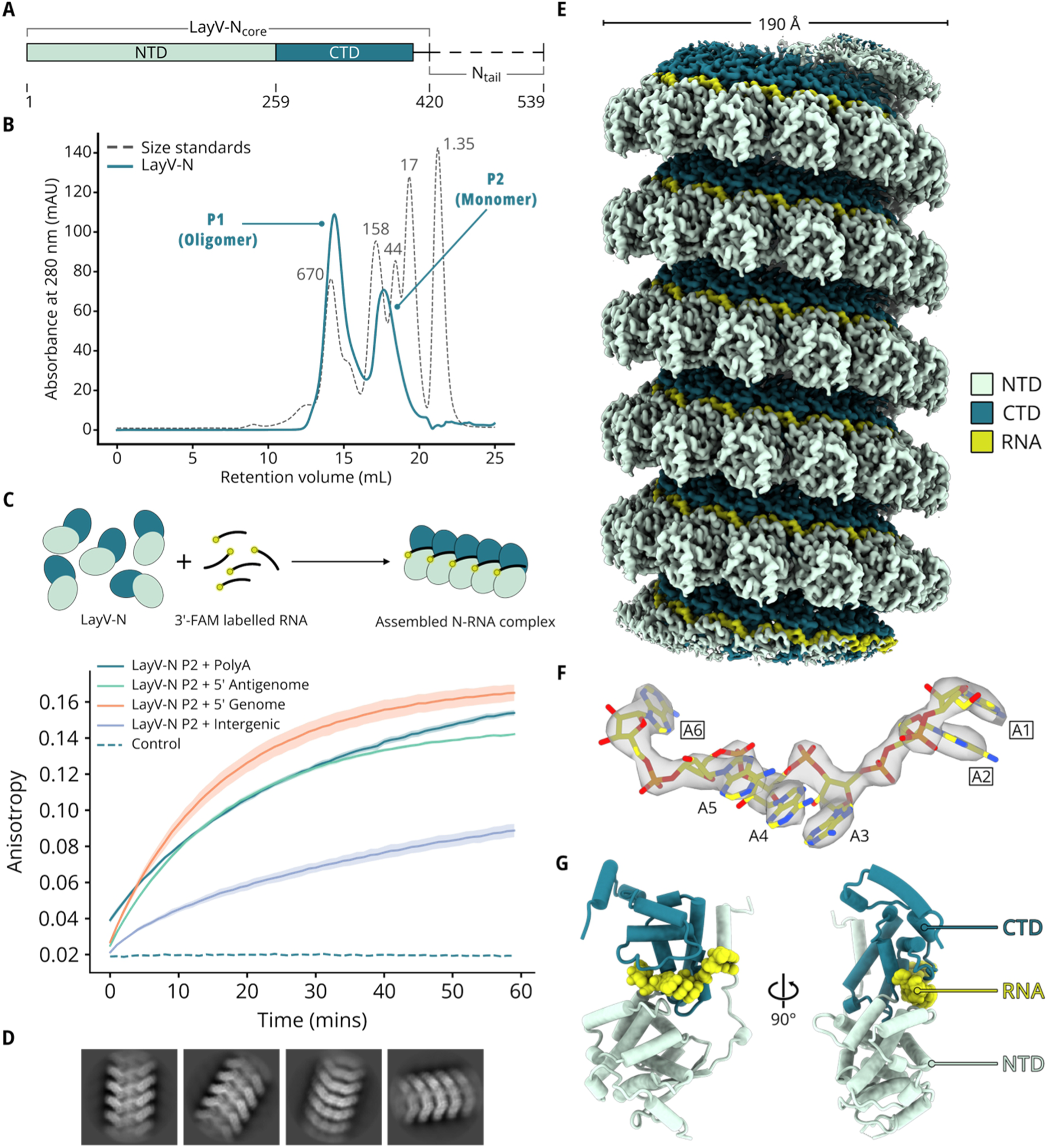
*in vitro* reconstitution of LayV-N_core_ nucleocapsid-like particles. **(A)** Schematic of full-length LayV-N highlighting the LayV-N_core_ encompassing the N- and C-terminal domains (NTD & CTD), and the N_tail_ with putative domain boundaries. **(B)** Representative size exclusion chromatogram of LayV-N_core_ purification with P1 and P2 populations labelled. The chromatogram is overlayed with that of size standards, the numbers in the standard curve are in kDa. **(C)** Schematic highlighting the *in vitro* reconstitution reaction for fluorescence anisotropy (top) and time-resolved anisotropy curves of reactions involving LayV-N_core_ P2 and different RNAs (bottom, see main text for RNA sequences). Standard deviations are plotted as uncertainty ranges along the curves. **(D)** Representative 2D class averages of the *in vitro* reconstituted nucleocapsid-like particles. **(E)** Final Coulomb potential map of the LayV-N_core_ + PolyA-RNA_6_ nucleocapsid-like particle coloured to indicate the NTD (green), CTD (teal), and RNA (yellow). Approximate diameter of the helix is shown. **(F)** Fit of the PolyA-RNA_6_ to its corresponding coulomb potential map with the nucleobases pointing into the helix (“three bases in”) highlighted by labels enclosed in rectangles. The adenines are numbered in the 5’ to 3’ direction. **(G)** Cartoon (cylinders) representation of the LayV-N-closed model bound to PolyA-RNA_6_ with same colour scheme as in panel F.

We expressed LayV-N_core_ in *E. coli* and purified it through immobilized metal affinity chromatography followed by size-exclusion chromatography (SEC). We observed two peaks (designated P1 and P2), consistent with the presence of two oligomeric states (Figure 1B). By comparison with a molecular mass standard, P1 corresponded to a species with an apparent mass exceeding 0.5 MDa, whereas P2 corresponded to a species below 80 kDa. Paramyxoviral N proteins expressed in *E. coli* often form assemblies composed of 12-14 protomers (*32–34*, *36–40*), suggesting that the first peak represents an assembled form of LayV-N_core_. Accordingly, we concluded that the second peak corresponds to the monomeric form, which is atypical due to the high propensity of N to oligomerize. To investigate the equilibrium between the two species, we collected P1 and P2 fractions, concentrated them, and re-analysed them by SEC. P1 fractions migrated reproducibly as a single oligomer peak (Supplementary Figure 1B), consistent with the notion that N assemblies are highly stable and resist dissociation (*51*). In contrast, concentrated P2 yielded two peaks, a large monomer peak at the P2 position, and a small oligomer peak at the P1 position (Supplementary Figure 1B). The observed behaviours indicated that the monomeric species irreversibly transitions into the oligomeric assembly, albeit with incomplete conversion. The ability to isolate predominantly non-oligomerized N proved serendipitous, as this required careful design of N-P fusion constructs (*33*, *34*, *44–46*, *52*) or co-expression with P constructs (*43*) in previous studies. Leveraging the monomeric LayV-N_core_, we decided to investigate RNA-uptake and assembly of henipaviral N.

### Monomeric LayV-N_core_ assembles into nucleocapsid-like particles upon incubation with RNA

To assess RNA uptake and assembly of LayV-N_core_ monomer fractions, we carried out time-resolved fluorescence anisotropy measurements with different sequences of fluorescently labelled RNA (Figure 1C). In line with previous observations from measles virus N (*52*), we observed sequence dependent differences. The efficiency of assembly was high for poly-A RNA, 5’-ends of genome (5’-ACCACACAAG-3’) or antigenome (5’-ACCAAACAAG-3’), while a pyrimidine-rich intergenic sequence (5’-CCUAAGUUUU-3’) showed low efficiency (Figure 1C). These observations are consistent with the notion of a rapid encapsidation of 5’-sequences as they emerge from the synthesizing polymerase during replication.

We then set out to structurally characterize the resulting RNA-bound assemblies. For this, we utilized LayV-N_core_ incubated with Poly-A RNA (PolyA-RNA_6_) overnight. The reaction was flash vitrified and subjected to cryogenic electron microscopy (cryo-EM). The resulting cryo-EM data revealed particles bearing the classic herringbone appearance of paramyxoviral nucleocapsids (*53*) (Supplementary figure 2C). Subsequent processing of this data led to 2D class averages which confirmed these particles to be helical assemblies (Figure 1D). Through helical 3D-reconstruction, we were able to resolve a Coulomb potential map of these helical assemblies to 3.1 Å resolution (using a gold standard FSC of 0.143) (Figure 1E).

The final map revealed that the helical assembly has a pitch of 56.6 Å with a left-handed twist of −27.9°, harbouring 12.92 subunits per turn. We could unambiguously assign high-resolution features of the LayV-N_core_ and the PolyA-RNA_6_ (Figures 1E and 1F). We refined a model into the map, resulting in a final structure of assembled LayV-N_core_ in a closed, RNA-bound conformation similar to related paramyxoviral N proteins (*32–40*) (Supplementary Figure 3). LayV-N-closed possesses a globular NTD and CTD holding the bound RNA in a cleft formed between the two domains (Figure 1G). Of the 420 residues of the protein, we were able model 397, along with the six adenine nucleotides of PolyA-RNA_6_. The non-modelled residues belong to a loop connecting the alpha helix α5, and beta sheet, β2, and the C-terminal extreme beyond residue 402 (Supplementary Figure 4). Each protomer in the helical assembly binds to one PolyA-RNA_6_ molecule with nucleotides oriented in the characteristic ‘three bases in, three bases out’ conformation, adhering to the rule of six (Figure 1F).

### Assembled LayV-N_core_ forms lateral contacts via the NT- and CT-arms and the insertion of an NTD loop

The helical assembly is organised by lateral contacts formed between two neighbouring protomers, similar to related paramyxoviral N assemblies (*32–40*) (Figures 2A, 2B). We analysed the assembly through PDBePISA (*54*) and found that the a surface area of ~3810 Å^2^ is buried at the interface between neighbouring protomers which is considerably larger than that of other paramyxoviruses for which N assemblies have been reported (Supplementary Table 1). The interaction of neighbouring protomers occurs through conserved mechanisms involving three regions of the N_core_ – an NTD loop comprising of residues 237-243, the NT-arm, and CT-arm. The NTD loop (237-243) of N_i_ protomer inserts itself into the so-called N-hole formed by the NT-arm and the NTD of N_i+1_ protomer (*39*) (Figure 2C). This loop insertion is facilitated by electrostatic interactions between the negatively charged residues (E312, D221, D95, D97) in the loops adjacent to the N-hole and positively charged residues R244 and K198 from N_i_. A similar NTD loop - N-hole interaction can be found in all other known paramyxoviral N assemblies (*32*, *33*, *35–40*) (Supplementary Figure 5), and it has also been observed in pneumoviruses (*55*, *56*) and filoviruses (*57–59*) (Supplementary Figure 6).

**Figure 2:**
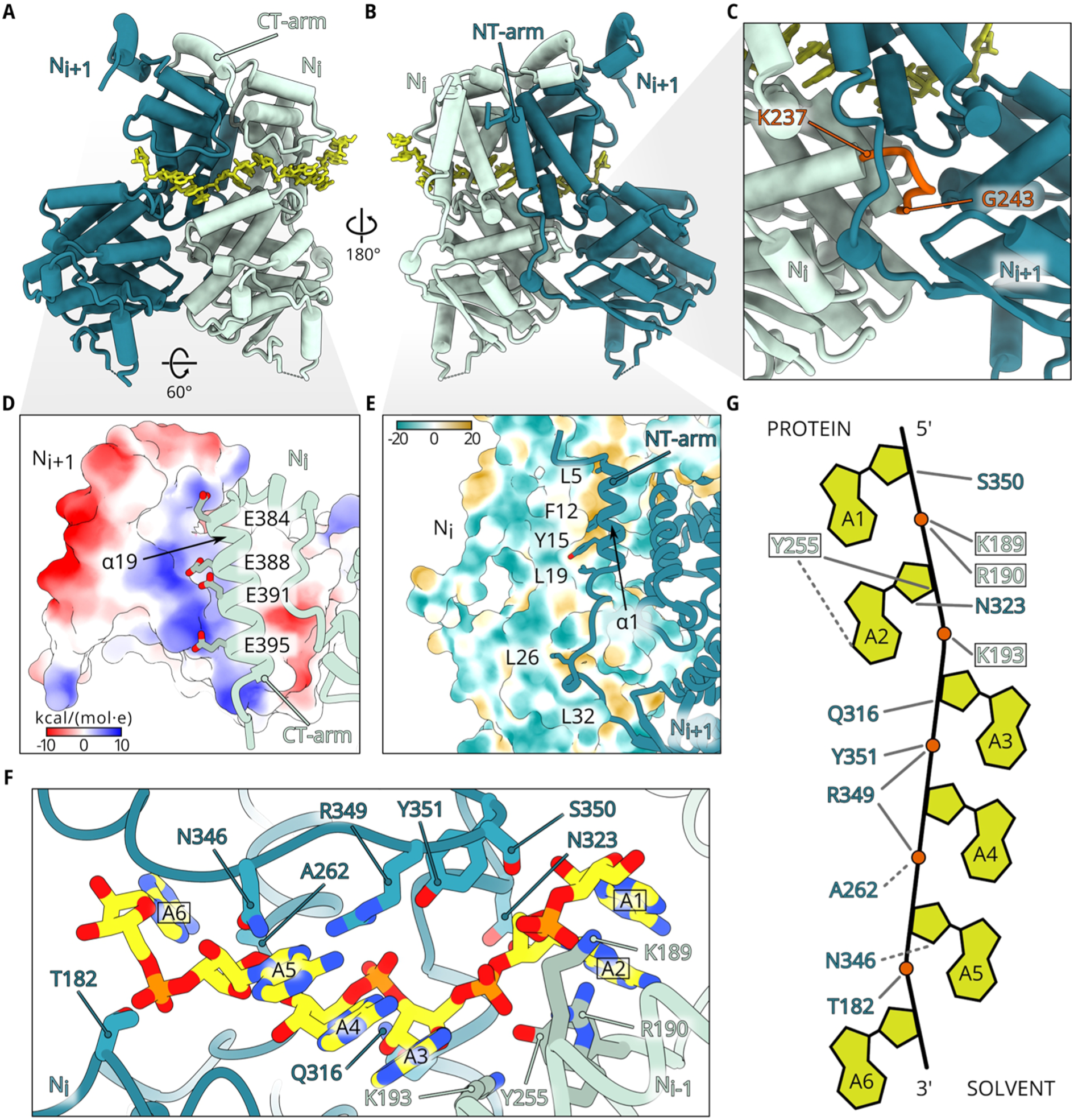
Lateral contacts and RNA-binding of the helical assemblies. **(A & B)** Cartoon (cylinders) representation of two neighbouring N_i_ and N_i+1_ LayV-N-closed protomers, coloured in teal and green, respectively. NT- and CT-arms are indicated and bound RNAs are shown as yellow sticks. **(C)** Closeup of the lateral interface focused on the insertion of NTD-loop (orange) into N-hole. Residue boundaries of the NTD loop are indicated. **(D)** Closeup of the interface between CT-arm of N_i_ (green ribbon) and the CTD of N_i+1_ (surface representation, coloured by electrostatic potential as indicated). Glutamates of the CT-arm are shown as sticks. **(E)** Closeup of the interface between NT-arm of N_i+1_ (teal ribbon) and the N_i_ protomer, shown as surface and coloured by hydrophobicity (brown = most hydrophobic, green = most hydrophilic). **(F)** Closeup showing the contacts between the PolyA-RNA_6_ (yellow sticks) and N_i_ (teal ribbon) and N_i+1_ (green ribbon) protomers. The residues that form hydrogen bonds with the RNA are labelled and shown as sticks in the respective colours of their protomers. The adenine nucleobases are labelled from 5’ to 3’ with the bases pointing towards the helix enclosed within rectangles. **(G)** Schematic showing the PolyA-RNA_6_ (nucleobases as yellow polygons and phosphates as orange spheres) and the hydrogen-bond forming residues from the N_i_ (teal) and N_i+1_ (green, enclosed with rectangles) protomers with the orientation of the nucleobases indicated as either ‘protein’ (bases in) and ‘solvent’ (bases out). Grey lines indicate hydrogen bonds formed via the sidechain atoms of the residues while dashed lines indicate hydrogen bonds formed through the main chain atoms.

The NT-arm and CT-arm of N_i_ protomer bind to N_i-1_ and N_i+1_ protomers, respectively (Figures 2A, 2B). The CT-arm comprises the alpha helix α19 and possesses four glutamate residues (E384, E388, E391, E395) that mediate electrostatic interactions with a positively charged patch in the CTD of N_i+1_ (Figure 2D). The NT-arm interaction with N_i-1_ is primarily hydrophobic in nature, mediated by residues L5, F12, Y15, and L19 from alpha helix α1 of N_i_ which bind to the hydrophobic groove located in the CTD of N_i-1_ (Figure 2E). The interaction of F12 with aromatic sidechains from N_i-1_ is well conserved in paramyxoviruses and has also been reported for NiV, HeV, and MeV (*32*, *36*, *40*) (Supplementary Figures 7A and 7B).

Apart from the lateral interface between two neighbouring protomers, another small interface between N_i-1_ and N_i+1_ protomers called the ‘elbow interface’ was recently reported for HeV (*40*). We could confirm the presence of a similar interface with ~96 Å^2^ of buried area formed by residues 3-6 in the N-terminal of N_i-1_ and residues 375, 378, and 379 of N_i+1_ (Supplementary Figure 7C).

### The PolyA-RNA_6_ is cradled in the positively charged cleft between NTD and CTD

Consistent with other reported closed N structures, the RNA in LayV-N-closed is bound in the cleft formed between the NTD and CTD. The PolyA-RNA_6_ is mainly held in place through its sugar-phosphate backbone which forms electrostatic interactions and hydrogen bonds with residues in the loop connecting alpha helices α12 and α13, and in alpha helix α7 (Supplementary Figure 8A and 8B). The RNA-binding cleft has a positively charged ‘floor’ through the residues K175, K189, R190, K193, K198 and R199, all but one (K175) of which are from α7 (Supplementary Figure 8B). Residues from N_i_ and N_i-1_ protomers both form hydrogen bonds with the sugar-phosphate backbone of one polyA-RNA_6_ molecule (Figure 2F). T182, Q316, N323, R349, S350, and Y351 of N_i_ protomer form hydrogen bonds through their sidechains while A262 and N346 form hydrogen bonds via their peptide backbone (Figure 2G). Four residues from N_i-1_, K189, R190, K193, and Y255, also form sidechain hydrogen bonds with the RNA backbone. Additionally, Y255 forms an additional hydrogen bond with N6 atom of the adenine nucleobase A2 through its mainchain O (Figure 2G). These non-covalent interactions cause the sugar-phosphate backbone of polyA-RNA_6_ to twist by ~180° every three nucleotides, thereby resulting in a ‘three bases in, three bases out’ orientation. The adenine nucleobases A1, A2, and A6 point into the helix whereas the base stack of the A3, A4, and A5 triad points towards the solvent (Figure 2G). The nucleobase A1 and A2 of the RNA bound to N_i_ stacks with A6 nucleobase of that bound to N_i-1_ to create the triad that points into the helix, leading to alternating triads that can be expected when a longer RNA is bound (Figure 2A).

The glutamine Q196 is well conserved across the paramyxoviruses (Supplementary Figure 8C), and in human parainfluenza virus 2, its equivalent glutamine (Q202) has been proposed to play a regulatory role in RNA synthesis ensuring that the rule of six is obeyed (*60*, *61*). In the MeV and PIV5, Q202 (the equivalent Q with the residue numbering of MeV and PIV5) binds RNA (*34*, *37*), but in LayV-N, Q196 is facing away from the nucleobases and does not seem to be directly involved in RNA binding (Supplementary Figure 8B).

### The helical assembly forms crucial vertical interactions via a conserved arginine of the CT-arm

While the lateral interfaces between adjacent protomers in N assemblies have been characterized for a number of paramyxoviruses, our understanding of the molecular determinants of vertical organization remains incomplete. This is due to the lack of well-resolved map density to identify side chain orientations at the vertical interface between protomers in previous studies (*32*, *34*, *35*, *39*). Our structure of LayV-N-closed is clearly resolved until residue 402 of the CT-arm, enabling us to assign sidechain positions (Figures 3A and 3B). Correspondingly, we see cryo-EM map density for sidechains of the NTD of N_i+13_ that point towards the CT-arm of N_i_, and we have identified those that could interact (Figure 3C).

**Figure 3:**
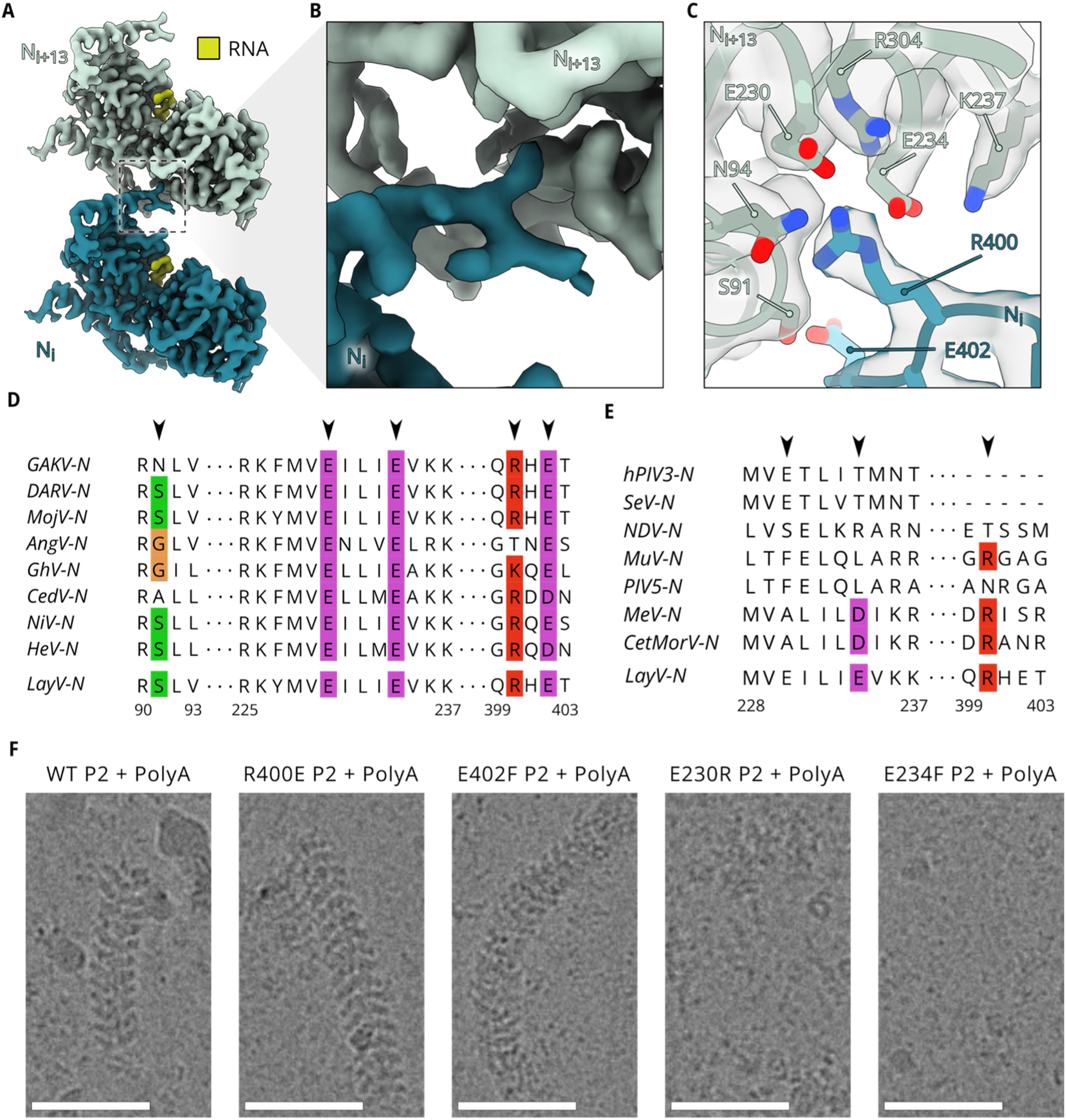
Vertical organization of the nucleocapsid-like particle. **(A)** Coulomb potential from the final map showing two protomers, N_i_ (teal) and N_i+13_ (green), vertically on top of each other in the helical assembly (RNA in both protomers highlighted in yellow). **(B)** Closeup view of the area indicated by the box in panel A to show the Coulomb potential map corresponding to the end of the CT-arm of N_i_. **(C)** Alternate view of the area shown in panel B, highlighting sidechain contacts, with the Coulomb potential shown with reduced opacity. The sidechains of residues present in the end of CT-arm of N_i_ (R400 and E402) and those of residues within hydrogen-bond forming distance to them from N_i+13_ are labelled and depicted as sticks. **(D)** Multiple sequence alignment of henipavirus N showing the conserved nature of residues 91, 230, 234, 400 and 402 (Black arrows; Residue numbering based on LayV-N). For clarity, only the regions of interest are shown. (GAKV = Gamak virus, DARV = Daeryong virus, AngV = Angavokely virus, GhV = Ghanaian bat virus, and CedV = Cedar virus). **(E)** Multiple sequence alignment of paramyxovirus N with black arrows indicating the equivalent residues to positions 230, 234, and 400 of LayV-N. For clarity, only the regions of interest are shown. (hPIV3 = Human parainfluenza virus 3, SeV = Sendai virus, NDV = Newcastle disease virus, MuV = mumps virus, PIV5 = Parainfluenza virus 5, MeV = Measles virus, and CetMorV = Cetacean morbillivirus). **(F)** Cryo-EM micrograph closeups of *in vitro* reconstitution reactions involving respective P2 populations of LayV-N WT and Mutants with PolyA-RNA_6_. Scale bars = 50 nm.

We could observe contacts between the protomers N_i_ and N_i+13_, specifically between R400 of N_i_ and a negatively charged pocket of N_i+13_ comprising S91, N94, E230, and E234. Based on the nature of these residues, it is likely that the assembly vertically interacts via electrostatic interactions and hydrogen bonds primarily anchored by R400 with S91, E230, and E234. The residue R400 is conserved, as part of the ‘QRHET’ motif found in the CT-arm of parahenipaviruses and as part of the ‘GR(Q/D)(D/E)’ motif of henipaviruses (Figure 3D). Similarly, the residues E230 and E234 are also conserved across all henipaviruses (Figure 3D). However, these three residues are not as well conserved within *Paramyxoviridae* (Figure 3E).

We then wanted to investigate whether the identified residues are vital in helical assembly formation. To this end, we purified LayV-N with point mutations E230R, E234F, R400E, and E402F and utilised the respective P2 fractions to be reconstituted with PolyA-RNA_6_. Samples of these reactions were flash vitrified and subjected to cryo-EM imaging. We observed normal helical assembly formation in the R400E and E402F mutants (Figure 3F). However, in both the E230R and the E234F mutants, assembly was severely disrupted, resulting in close to no intact helical filaments being observable (Figure 3F). These observations indicate that interactions involving residues E230 and E234 play an important role in mediating the vertical organization. The conservation of contact residues in this region suggests that the nucleocapsids of other henipaviruses are vertically stabilized utilizing equivalent interactions as in LayV.

### LayV-N_core_ can form an RNA-free ring-assemblies that transition into helical structures upon RNA incubation

Next, we set out to characterize the large oligomer fractions of P1 of our LayV-N_core_ purification (Figure 1B). Cryo-EM data revealed that this sample contained ring-like assemblies of LayV-N_core_ comprising 13 protomers. Further processing of rings led to a Coulomb potential map of 2.61 Å resolution (Figure 4A). The ring-like assembly possessed a diameter of about 195 Å and contained density for 13 protomers. Unexpectedly, and opposed to previous reports of ring-assemblies (*35*, *37*), the cryo-EM map was lacking any density for host-derived RNA (Figure 4A). The absence of RNA was corroborated by a A_260_/A_280_ measurement of 0.75. The lack of bound RNA is in stark contrast to all previous recombinant N ring assemblies in which N has taken up random host RNA upon assembly.

**Figure 4:**
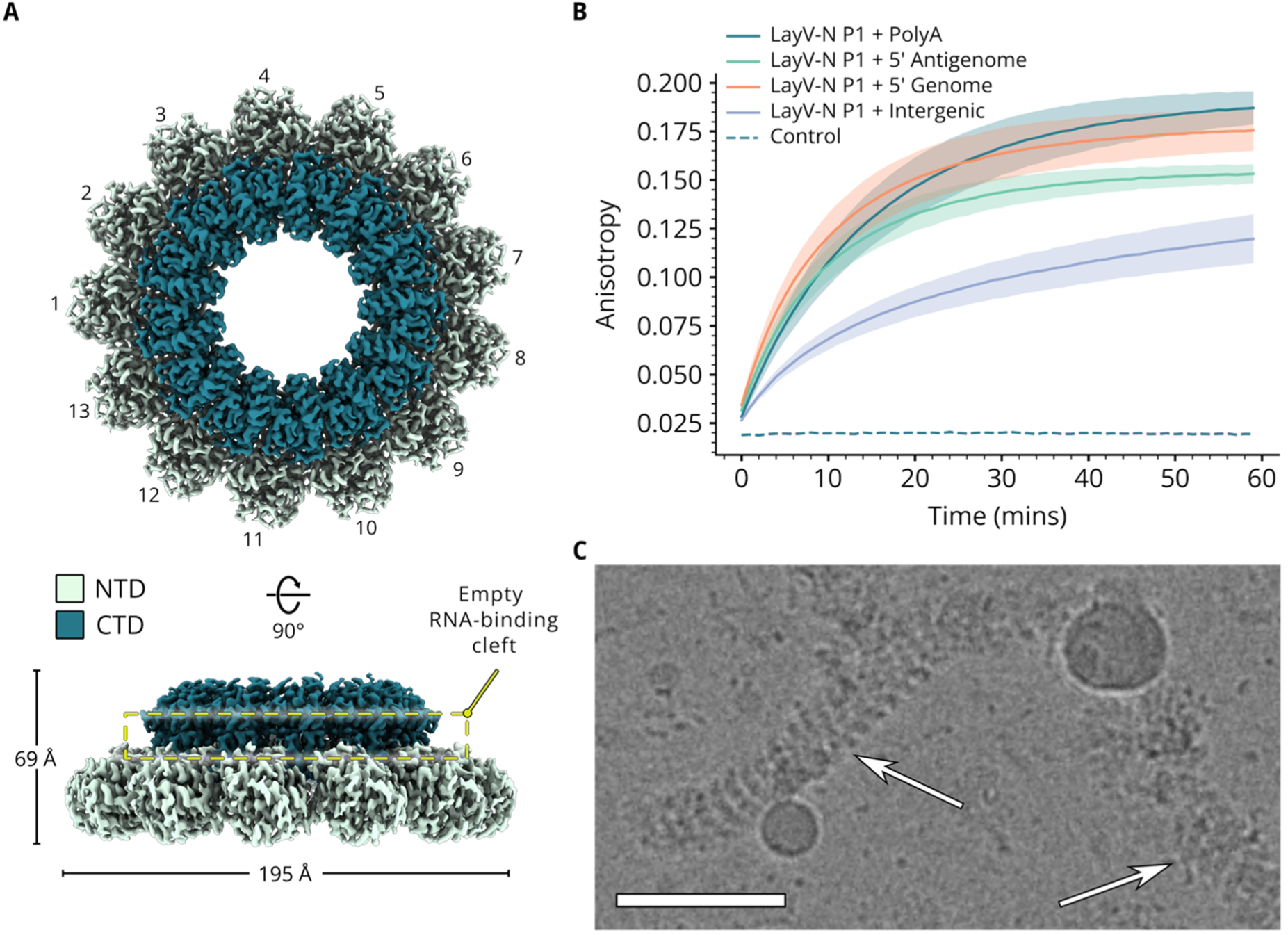
RNA-free LayV-N_core_ assembly and its ring-like organization. **(A)** Top and side views of the cryo-EM map of the LayV-N_core_ ring assembly with features corresponding to 13 protomers indicated. Map is coloured to indicate the NTD (green) and CTD (teal). Also indicated in the sideview is the empty RNA-binding cleft. **(B)** Plot of the anisotropy curves of reactions involving LayV-N P1 and different RNAs (see main text for RNA sequences). Standard deviations are plotted as uncertainty ranges along the curves. **(C)** Cryo-EM micrograph closeup of *in vitro* reconstituted LayV-N WT P1 with PolyA-RNA_6_. Arrows highlighting the helical assemblies. Scale bar = 50 nm.

To assess whether LayV-N rings can convert to helical assemblies, we incubated them with the same fluorescently labelled RNA sequences as described earlier for LayV-N_core_ P2 and carried out time-resolved fluorescence anisotropy measurements. We observed comparable trends to those of the monomer fraction, with a lower assembly efficiency using pyrimidine-rich intergenic RNA than with A-rich sequences (Figure 4B). To validate whether the observed higher-order assemblies were helical, we flash vitrified an overnight reaction of LayV-N rings incubated with PolyA-RNA_6_ and screened the sample using cryo-EM. The resulting cryo-EM images contained many helical filaments bearing the characteristic herringbone pattern, confirming that the RNA-free rings can accommodate the PolyA-RNA_6_ and transition to form helical assemblies (Figure 4C).

### LayV-N_core_ can adopt open and closed conformations in the assembled state

To further improve the map of ring-like assemblies, we performed symmetry expansion followed by local refinement centred on a single protomer (Figure 5A). Refinement of a model into the map density revealed a conformational change of LayV-N_core_ compared to the RNA-bound assembly, with the CTD moved further away from the NTD, opening the RNA-binding cleft (Figure 5B).

**Figure 5:**
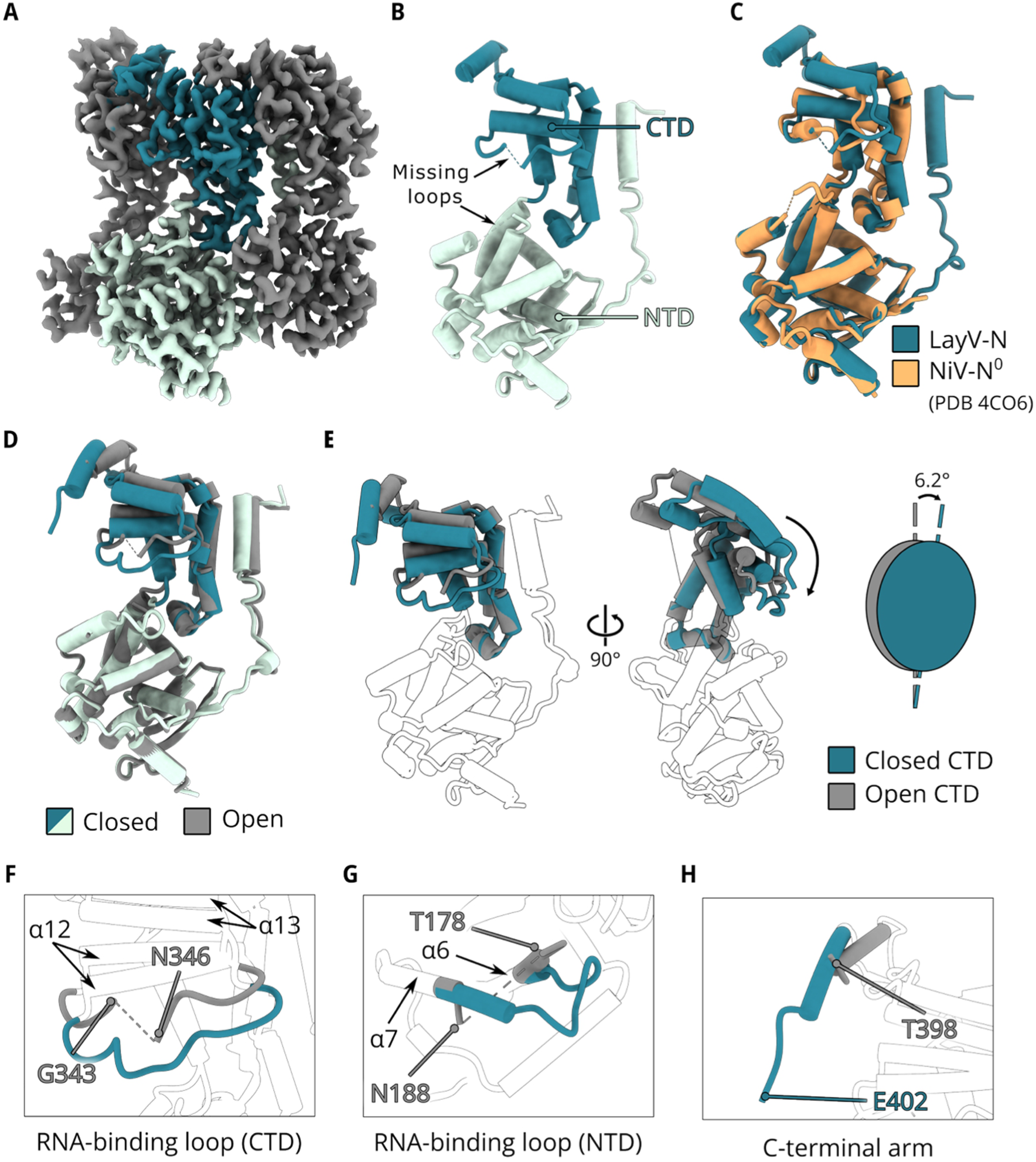
Comparison of the open and closed LayV-N models. **(A)** Coulomb potential map of the local refined region focusing on one protomer which is shown in green and teal (NTD and CTD, respectively). **(B)** Cartoon (cylinders) representation of the LayV-N-opened model coloured in the same scheme as earlier to indicate NTD and CTD. Missing RNA-binding loops are indicated. **(C)** Structural alignment of LayV-N-open with the NiV-N^0^-P model (PDB 4CO6). The alignment yielded an RMSD of 0.95 Å. Alignment carried out in UCSF ChimeraX 1.7.1 (*62*). **(D)** Alignment of LayV-N-closed (NTD in green and CTD in teal) with LayV-N-open (grey) models. **(E)** Panel highlighting the difference in orientation of the CTDs between the two models using two different views of the superimposition from panel A. Here, only the CTDs are coloured (using the same colour scheme as panel A) with the superimposed NTDs made transparent. A schematic representation of the relative difference in orientation, indicating the movement of 6.2° of the CTDs, is also shown. **(F, G, and H)** Panels highlighting the difference in RNA-binding loops of CTD and NTD between the open and closed models, and the visualisation of the extension of CT-arm in the closed model compared to the open model. In all three panels, the open model is coloured in grey, and the closed model is coloured in teal.

Structural alignment of the model with that of open NiV-N structure (NiV N^0^-P, PDB 4CO6) (*43*), yielded an RMSD value of 0.95 Å, validating that this LayV-N_core_ is indeed in an open conformation, and we will refer to it as LayV-N-open (Figure 5C and Supplementary Figure 9). We were able to model 381 amino acids of LayV-N-open, with small portions missing in the RNA binding loops and in the C-terminus, presumably due to flexibility (Figure 5B). The rings are organised by lateral contacts, analogous to those described earlier for the helical assembly (Supplementary Figure 10).

We compared the structures of LayV-N-closed and LayV-N-open by structural alignment (Figure 5D). The comparison of the two structures highlighted the key differences between the conformations that are primarily imposed by the presence (or absence) of RNA. First, we noticed that the CTD of the LayV-N-closed is seen clamping down towards the NTD to bind the RNA, moving 6.2° downwards with respect to the centre of mass of NTD (Figure 5E). This observed movement of CTD is what distinguishes the closed conformation from the open conformation. Secondly, we also observe changes in the RNA-binding loops of NTD and CTD (Figures 5F and 5G). In the open conformation, map density of residues A344 and L345 in the CTD loop between alpha helices α12 and α13 was disordered. Similarly, the residues 179-187 could not be modelled in the loop connecting alpha helices α6 and α7 highlighting the flexible nature of these loops. However, these disordered regions become ordered in the presence of RNA in the closed conformation indicating a folding upon binding mechanism.

Lastly, we could observe a clearer Coulomb potential map of residues 399-402 in the CT-arm of helically assembled LayV-N-closed compared to the protein in the ring-like assembly. This is likely due to vertical stabilization from protomers in the consecutive turn of the helical assembly, enabling their visualization (Figure 5H).

## Discussion

Nucleoproteins are by far the most abundant viral protein produced in cells infected by paramyxoviruses (*63*). By encapsidating the viral RNA genome to form the nucleocapsid as protection against immune recognition (*64*) and host nucleases (*65*), and by interacting with the polymerase complex, nucleoproteins constitute a key component of the RNA synthesis machinery of these viruses. During both transcription and replication, the polymerase complex initiates *de novo* RNA synthesis utilising the nucleocapsid as the template (*66*, *67*). The polymerase complex thus needs to read the RNA packaged by the nucleoprotein polymer to generate a complementary product. However, the mechanism by which the polymerase complex is able to access and read the fully occluded RNA sequence in the nucleocapsid is unclear.

Here we characterise cryo-EM structures of assembled LayV-N_core_ in both its open and closed states. Obtaining these structures was the result of serendipitous, atypical behaviour of LayV-N_core_ in solution, where we find that it co-exists in two populations with appreciable abundance – as an RNA-free monomer and RNA-free oligomer. This is in contrast to previous studies on MeV (*34*) and CeMV (*33*), where to stabilize an RNA-free monomeric N, it needed to be covalently fused to a region of P. It may very well be that in LayV-N the conformational equilibrium is shifted further towards an RNA-free state as compared to other paramyxoviruses, enabling our study. We further show that the monomeric LayV-N_core_ readily encapsidates supplied RNA, leading to the formation of a nucleocapsid-like helical assembly, with LayV-N_core_ in a closed conformation, biting down on the supplied RNA molecule. As in other paramyxoviruses, the assembly is stabilized laterally by contacts mediated by the NT-arm, CT-arm, and the NTD loop (residues 237-243). However, our structure additionally resolves an interface involved in maintaining contact between N protomers of consecutive turns of the helical assembly. Specifically, we identified a conserved arginine residue at position 400 of the CT-arm, which contacts two conserved glutamates of the vertically neighbouring protomer. Site-directed mutagenesis demonstrates that the glutamates are crucial for the formation of helical assemblies, providing new insight into the molecular keystones enabling nucleocapsid construction. We also demonstrate that the RNA encapsidation by monomeric LayV-N_core_ is sequence dependent with 5’ genomic and antigenomic sequences being rapidly packaged compared to pyrimidine-rich intergenic RNA. This finding is consistent with what has previously been reported for MeV-N (*52*).

Notably, we were also able to characterize at high-resolution, LayV-N in an RNA-free assembled state in absence of the P protein. Ring-like assemblies are not uncommon with single rings comprised of 13 protomers reported for PIV5 (*37*) and MuV (*35*). Beyond *Paramyxoviridae*, RNA bound ring-like assemblies have also been observed for *Pneumoviridae* (*68*, *69*), *Rhabdoviridae* (*70*, *71*), and *Bornaviridae* (*72*). It is unclear what might be the significance of these ring-like assemblies in viral infections, however they are also present *in situ* within virions (*73*). It has been suggested that these rings might bind the short leader RNA transcripts produced by the polymerase complex during the initial phase of transcription (*74*, *75*) or might serve as a temporary source of metastable N readily available to be recruited by P (*40*). We observe that RNA-free rings can undergo a transition to helical assemblies in presence of supplied RNA. This adds weight to the idea that rings could serve as a storage form of N that can be utilized as building blocks for nucleocapsids when the need arises. Future cryo-ET studies of infected cells may shed more light on whether N-rings are present *in situ* and what their functions might be.

To date, all reported paramyxoviral N proteins in the RNA-free open state were in presence of the P_NTD_ (*43–46*). In contrast, assemblies have been either bound to RNA of the expression host (*32*, *35–40*) or to added synthetic RNA (*33*, *34*). The structures of these assemblies demonstrated that N adopts the closed state when bound to RNA. This led to the notion that N assembly, RNA-binding, and the conformational switch from open to closed state are coupled. It is thought that RNA will have a stabilizing effect on the closed conformation, similar to that elicited by the presence of P_NTD_ for the open state, while also enabling oligomerisation via lateral contacts (*76*). Our high-resolution characterization of RNA-free N rings in absence of P_NTD_ demonstrates that N can be stable within an oligomeric context and without RNA, painting a more nuanced picture.

Our data provides structural evidence suggesting that assembled N is capable of releasing RNA, independent of P_NTD_, and without disrupting the assembly, thereby providing a pathway to relinquish the RNA-template to the synthesizing polymerase complex. While the reason that we are able to capture this structural snapshot may very well be due to the unique equilibrium position of LayV-N, it may be indicative of the general behaviour of paramyxovirus nucleoproteins. Local RNA release, without nucleocapsid disassembly, is compatible with previous hypotheses stating that a conformational change in the α7 helix adjacent to the NTD RNA-binding loop, is sufficient to release the RNA from the nucleocapsid (*77*). Disengagement of the RNA-binding loop might be triggered by the attachment of the P_XD_-N_αMoRE_ helical bundle to N (*49*), in the context of the polymerase complex traversing the nucleocapsid.

## Materials and Methods

### LayV-N_core_ construct design

We retrieved the full-length coding sequence of LayV-N from NCBI Genbank (accession number: OM101125.1) and predicted the disorder score for each residue in this sequence using Metapredict v3 (*50*). Based on the disorder score plot, we designed an expression construct for LayV-N_core_ (residues 1-420) included a 6xHistidine tag, V5 tag and a tobacco etch virus (TEV) protease cleavage site upstream of the coding sequence. We ordered this construct as a gene insert, codon-optimized for *Escherichia coli*, pre-cloned into pET151/D-TOPO expression vector, from GeneArt (ThermoFisher Scientific). We subsequently confirmed the sequence of the construct through Sanger sequencing.

Point mutant constructs of LayV-N_core_ (E230R, E234F, R400E, and E402F) were designed and obtained similarly to the wild type.

### Protein expression and purification

We expressed and purified all proteins involved in this study as follows unless stated otherwise. We transformed *E. coli* BL21 (DE3) pLysS competent cells with the LayV-N_core_ encoding plasmid and expressed the transformants in Luria Bertani media (Invitrogen), containing 50 μg/mL carbenicillin and 34 μg/mL chloramphenicol. We started by culturing the cells at 37°C while shaking (170 RPM) until OD_600_ measurements reached 0.5. We then cooled the cultures to 16°C and induced recombinant protein expression by adding Isopropyl β-D-thiogalactopyranoside (IPTG) to a final concentration of 0.4 mM. Post-induction, we continued culturing at 16°C overnight (16-18 hours) while shaking (170 RPM). We then followed a purification protocol based on the one described by Ker *et al.*, for NiV-N (*36*). Briefly, we harvested the cells by centrifuging (4500 x *g*, 4°C) for 20 minutes and resuspended the pelleted cells in 50 mL of lysis buffer (20 mM Tris-HCl, pH 7.5, 1 M NaCl, 1 M Urea, and 10% (v/v) glycerol, supplemented with a Pierce™ protease inhibitor tablet (ThermoFisher Scientific)) per L of culture. We lysed the resuspended cells using sonication (40% amplitude for 20 cycles of 10 s on, 30 s of) and then clarified the resulting lysate by centrifugation (48000 x *g*, 4°C) for 45 minutes.

To purify the recombinant LayV-N_core_ proteins, we employed immobilized metal ion affinity chromatography. First, we applied the clarified lysate to a gravity flow column (BioRad) packed with 1 mL of HisPur™ Ni-NTA superflow agarose (ThermoFisher Scientific) that was pre-equilibrated with the binding buffer (20 mM Tris-HCl, pH 7.5, 1 M NaCl, and 10% (v/v) glycerol). We subsequently washed the column with 20 column volumes (CV) of binding buffer followed by 10 CV of binding buffer containing 35 mM imidazole. We then eluted LayV-N_core_ proteins using a stepwise gradient of imidazole diluted in binding buffer. Our elution steps involved passing 5 CV of binding buffer supplemented with 70 mM, 150 mM, and 300 mM of imidazole through the column and we collected flow-through at every step for SDS-PAGE analysis. We pooled the eluted fractions containing higher amounts of LayV-N_core_ and performed another purification step with size-exclusion chromatography (SEC). With SEC, we purified the pooled sample using a Superose 6 increase 10/300 GL column (Cytiva) that was pre-equilibrated in the SEC buffer (20 mM Tris-HCl, pH 7.5, 500 mM NaCl). Post SEC, we processed the fractions corresponding to P1 and P2 individually. We concentrated P1 and P2 samples using 100 kDa MWCO and 30 kDa MWCO Amicon Ultra centrifugal filter units, respectively (Merck) and stored them at 8°C until further use.

### Time-resolved fluorescence anisotropy

We identified the viral RNA sequences corresponding to the 5’ ends of both genome and antigenome from the NCBI GenBank deposition of LayV strain Cl17-6 (accession number: PQ641425.1). For intergenic RNA, we used the sequence located between the N and P ORFs from the same deposition. We ordered the identified sequences and a polyadenine sequence as synthetic RNA 10mers, labelled at the 3’ end with 6-FAM from Merck. The ordered sequences are as follows: PolyA-RNA_10_ (5’ AAA AAA AAAA 3’), 5’ Genome-RNA_10_ (5’ ACC ACA CAAG 3’), 5’ Antigenome-RNA_10_ (5’ ACC AAA CAAG 3’), and Intergenic-RNA_10_ (5’ CCU AAG UUUU 3’).

For time-resolved fluorescence anisotropy experiments, we mixed purified LayV-N_core_ (2.5 µM) with the respective RNA in a 5:1 (Protein:RNA) molar ratio to a final reaction volume of 40 µL with SEC buffer (20 mM Tris-HCl, pH 7.5, 500 mM NaCl). We prepared the reactions in black 384-well microplates (BRANDTECH scientific) and measured the emission fluorescence intensities parallel (F||) and perpendicular (F ⊥) to the excitation light plane using a CLARIOstar Plus plate reader (BMG Labtech). Measurements were made every minute using excitation/emission wavelengths of 482/530 nm, with 50 flashes per well per cycle, for a total reaction time of 60 minutes. The plate was shaken at 300 RPM between each measurement. For each reaction, the anisotropy (r) was calculated based on the following expression:

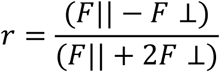

To present the anisotropy data, we averaged measurements from three technical replicates for each biological replicate, and we plotted the mean from three such biological replicates for each reaction. We calculated the error bars as mean ± standard deviation using the values obtained for the biological replicates and showed them as uncertainty ranges accompanying the curves.

### *In vitro* reconstitution of nucleocapsid-like particle

To reconstitute nucleocapsid-like particles for cryo-EM, we mixed monomeric LayV-N_core_ (10 μM) from the concentrated P2 sample with 5-fold molar excess of synthetic polyA-RNA_6_ (5’ AAA AAA 3’) (Eurofins Genomics). We diluted the reaction to a final buffer composition of 20 mM Tris-HCl (pH 7.5) and 200 mM NaCl, and incubated them at 8°C overnight (16-18 hours) before use.

### Cryo-EM sample preparation and data acquisition

To prepare cryo-EM samples, we applied 3 μL each of the *in vitro* reconstituted samples (WT and mutants) and the oligomeric LayV-N_core_ P1 (0.5 mg/mL) to glow-discharged Quantifoil R 2/2 Cu 300 and R 2/1 Cu 300 holey carbon grids, respectively (Quantifoil Micro Tools). We then blotted the grids for 5 s before vitrifying them in liquid ethane using a Vitrobot mark IV robot (FEI) operated at room temperature with 100% humidity.

We acquired cryo-EM data from both the *in vitro* reconstituted sample and the oligomeric LayV-N_core_ P1 using a 300 keV Titan Krios cryo-transmission electron microscope (FEI) equipped with a Falcon 4i detector and a Selectris energy filter (ThermoFisher), located at the Umeå Centre for Electron Microscopy (UCEM). We automated data acquisition using EPU software v3.7.0 (ThermoFisher Scientific) and collected 14598 and 21141 movies for the two samples respectively. Additional data acquisition parameters can be found listed in Supplementary Table 2.

We collected cryo-EM images for *in vitro* reconstitutions of LayV-N_core_ WT and mutants with a 200 keV Glacios cryo-transmission electron microscope (ThermoFisher Scientific) equipped with a Falcon 4i detector, located at UCEM. We imaged the reconstituted samples using a total electron dose of 51.89 e^-^/Å^2^ (at an exposure rate of 9.58 e^-^/px/s for 4.92 s), at a magnification of 150 000x which yields a nominal pixel size of 1.05 Å/px.

### Image processing

We processed all cryo-EM data using CryoSPARC software platform v4.5.3 (*78*) unless stated otherwise.

#### In vitro reconstituted nucleocapsid-like particles

We started by aligning the movie frames and estimating their contrast transfer functions (CTF) using the patch motion correction and patch CTF estimation jobs. Following manual curation of the motion-corrected micrographs, where we discarded the ones with thick ice, bad CTF fits, and poor defocus, we used the blob picker to pick ~4 million particles.

We inspected this initial particle set and removed duplicates and other suboptimal picks to obtain a curated particle set of ~2 million particles, out of which 1 546 278 were extracted, Fourier-cropped to a pixel size of 5.24 Å/px. We performed 2D classification on the extracted particles and selected ten of the resulting classes to generate templates for template-based picking. From the ~4 million particles picked by the template picker, we extracted 1 430 740 particles, Fourier-cropped to a pixel size of 5.24 Å/px. We then performed a 2D classification of the extracted particles and selected nine classes, containing a total of 226 801 particles for ab-initio reconstruction. We reconstructed four volumes from which the best one was chosen as the initial volume for 3D refinement with C1 symmetry involving the selected 226 801 particles.

After estimating the helical symmetry parameters using the newly refined volume, we extracted 225 614 uncropped particles from the particle set utilized for helical refinement. We performed another round of ab-initio reconstruction followed by helical refinement which ended up utilizing 170 777 uncropped particles. This was followed by an additional round of helical refinement with the same 170 777 particles, after imposing the estimated helical symmetry parameters from earlier. We thus obtained the final map of the *in vitro* reconstituted helical assembly at 3.1 Å with a helical rise and twist of 4.4 Å and −27.9°, respectively.

#### RNA-free LayV-N_core_ rings

Following the same initial approach as described above, we picked ~3.7 million particles from 19 251 curated micrographs of the oligomeric LayV-N_core_ dataset. We then extracted about 3 million of the picked particles, Fourier-cropped to a pixel size of 5.24 Å/px and subjected the extracted particles to a round of 2D classification. We observed that 36% of the classified particles (~1 million) resembled single rings.

We performed ab initio reconstruction on the particle set obtained from the 2D classes of rings followed by non-uniform refinement, to achieve an initial volume at 11.3 Å. We then extracted 925k particles from the particle set of this refinement job from un-binned micrographs and performed another non-uniform refinement by imposing a C13 symmetry. Using the resulting mask, we classified the particles in 3D and took forward the 541 364 particles that were classified into the three best looking classes for another round of non-uniform refinement with C13 symmetry. This gave us a refined volume at 2.6 Å but the NTD suffered from some anisotropy, perhaps due to a lack of side views, particularly in the region around α5. To improve the map quality, we set out to locally refine the volume focusing on a single protomer. We started by creating a density map encompassing a protomer using the ‘molmap’ command in UCSF Chimera 1.17.3 (*79*). We then created a monomer-mask from this map with the ‘Mask Creation’ job in RELION 3.1.0 (*80*), defining the region to be locally refined in the ring. Lastly, we ran a local refinement job utilising the mask and a particle set of ~7 million obtained through symmetry expansion to achieve a final map of a single protomer at 2.5 Å. The resulting map had greatly improved quality (see Figure 5A) and was used for model building.

### Model building, and refinement

We built the model of both our structures starting by rigid body fitting a model of LayV-N_core_ predicted using ColabFold (*81*), a structure prediction software which utilizes AlphaFold2 (*82*). We then iteratively refined the model using COOT (*83*) and the Real-space refinement job (*84*) in Phenix v1.21.1 (*85*). After 20 and 9 rounds of real-space refinement for LayV-N-closed and LayV-N-open, respectively, we obtained the final models presented here. Model refinement and validation statistics can be found in Supplementary Table 2.

### Data visualization

We analysed the structural interfaces using PDBePISA (‘Protein interfaces, surfaces and assemblies’ service PISA at the European Bioinformatics Institute) (http://www.ebi.ac.uk/pdbe/prot_int/pistart.html) (*54*). We used UCSF Chimera 1.17.3 (*79*) and UCSF ChimeraX 1.7.1 (*62*) to visualize all the structural data shown here. We performed multiple sequence alignments with Clustal Omega (*86*, *87*) and visualized them with clustal X colour scheme using Jalview v2.11.5.0 (*88*). We obtained the protein sequences for alignment from NCBI GenBank using the following accession numbers: LayV (OM101125.1), HeV (NC_001906.3), NiV (NC_002728.1), AngV (ON613535.1), CedV (NC_025351.1), GhV (NC_025256.1), GAKV (MZ574407.1), MojV (NC_025352.1), DARV (MZ574409.1), MeV (NC_001498.1), CetMorV (NC_005283.1), MuV (NC_002200.1), PIV5 (NC_006430.1), NDV (NC_075404.1), hPIV3 (NC_001796.2), SeV (NC_075392.1). We utilized ggtree (*89*) to obtain the phylogenetic tree shown in Supplementary Figure 1 using the amino acid sequences of the respective L proteins. We used the matplotlib package for python (v3.9.0)(*90*) to plot chromatograms, disorder scores, and time-resolved fluorescence anisotropy data utilising colours from ColorBrewer2.0 (*91*). Figure panels were compiled using Inkscape v1.3.2.

## Acknowledgements

We are grateful for financial support from the Kempe Foundation (grants JCSMK22-0127 and JCK 23135), the Umeå Centre for Microbial Research, and Pandemifonden. We thank Irina Gutsche (Institut de Biologie Structurale, Grenoble, France) for scientific discussions. We are also grateful for support from UCEM staff, especially Lorène Gonnin, Michael Hall, Sara Sandin, and Erin Schexnaydre for assistance with cryo-EM data acquisition. We also thank members of the Renner and Carlson labs for their scientific input and discussions.

Cryo-EM data were collected at the Umeå Centre for Electron Microscopy (UCEM), a node of the Cryo-EM Swedish National Facility, funded by the Knut and Alice Wallenberg, Family Erling Persson and Kempe Foundations, SciLifeLab, Stockholm University and Umeå University.

We gratefully acknowledge the support from the Chemical Biology Consortium Sweden (CBCS), a national research infrastructure funded by the Swedish Research Council (dr.nr.2021-00179), SciLifeLab, and its hosting Universities.

## Author contributions

CRediT authorship contribution statement

Rupesh Balaji Jayachandran: Conceptualization, Investigation, Visualization, Methodology, Formal analysis, Validation, Writing - original draft, review & editing.

Erwan Quignon: Investigation, Visualization, Methodology, Formal analysis, Validation, Writing – original draft, review & editing.

Max Renner: Conceptualization, Supervision, Investigation, Formal analysis, Funding acquisition, Resources, Writing - original draft, review & editing.

## Data availability

The coordinates and density maps generated in this study have been deposited in the Protein Data Bank (PDB) and the Electron Microscopy Databank (EMDB), respectively. They can be accessed using the following accession codes: LayV-N-closed (PDB XXXX), LayV-N-Open (PDB XXXX), *in vitro* reconstituted helical assembly of LayV-N_core_ + PolyA-RNA_6_ (EMD-XXXXX), RNA-free LayV-N_core_ ring assembly (EMD-XXXXX), and the locally refined map of a single protomer from LayV-N_core_ ring (EMD-XXXXX). Raw data underlying figures 1, 3, 4, and 5 are provided in the Source Data files.

