## Supplementary Data for "Open and closed forms of assembled henipavirus nucleoprotein suggest structural basis of genome access"

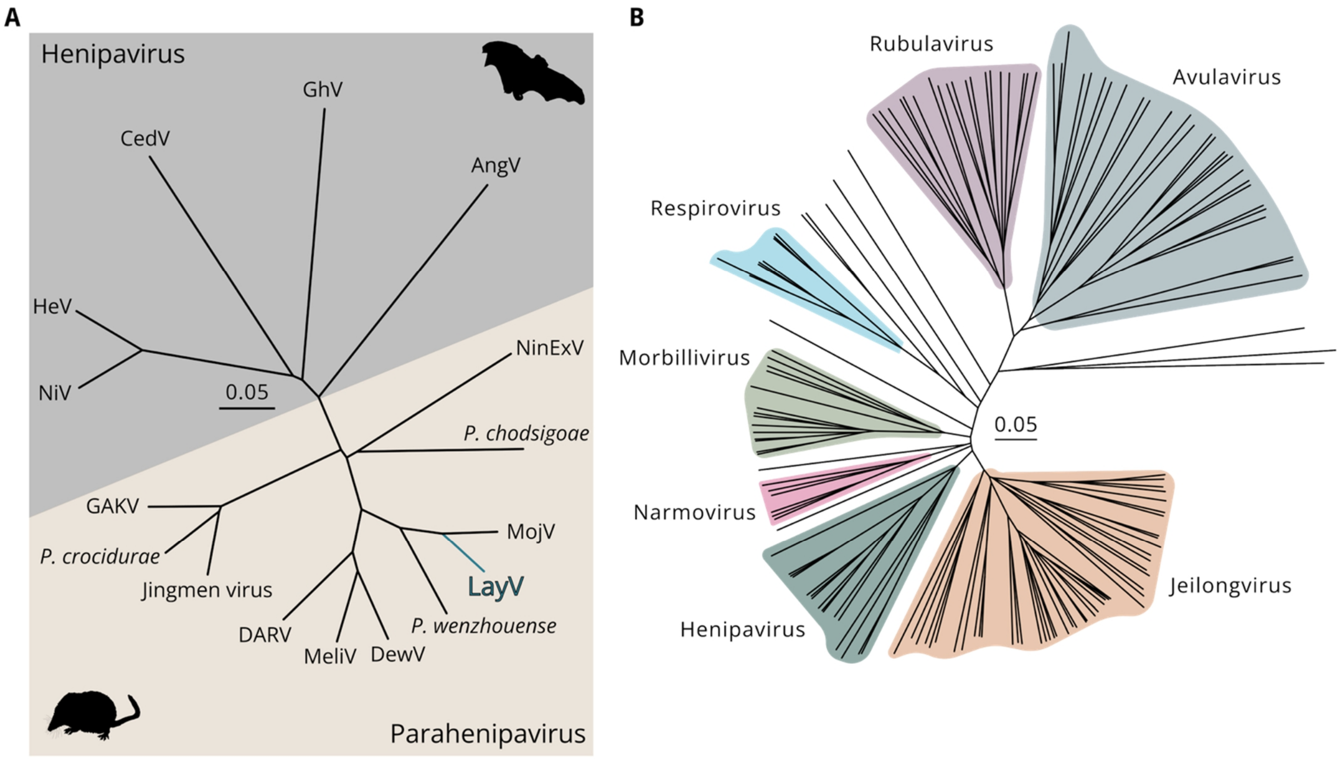


**Supplementary Figure 1: Phylogeny of henipaviruses and paramyxoviruses. (A)** Unrooted phylogenetic tree of the *Henipavirus* and *Parahenipaviruses* genera. (NiV = Nipah virus, HeV = Hendra. Virus, CedV = Cedar virus, GhV = Ghanaian bat virus, AngV = Angavokely virus, NinExV = Ninorex virus, MojV = Mòjiāng virus, DewV = Denwin virus, MeliV = Melian virus, DARV = Daeryong virus, and GAKV = Gamak virus). **(B)** Unrooted phylogenetic tree of the *Paramyxoviridae* family, with major subfamilies coloured differently. Both phylogenetic trees were generated using the amino acid sequences of the respective L proteins.

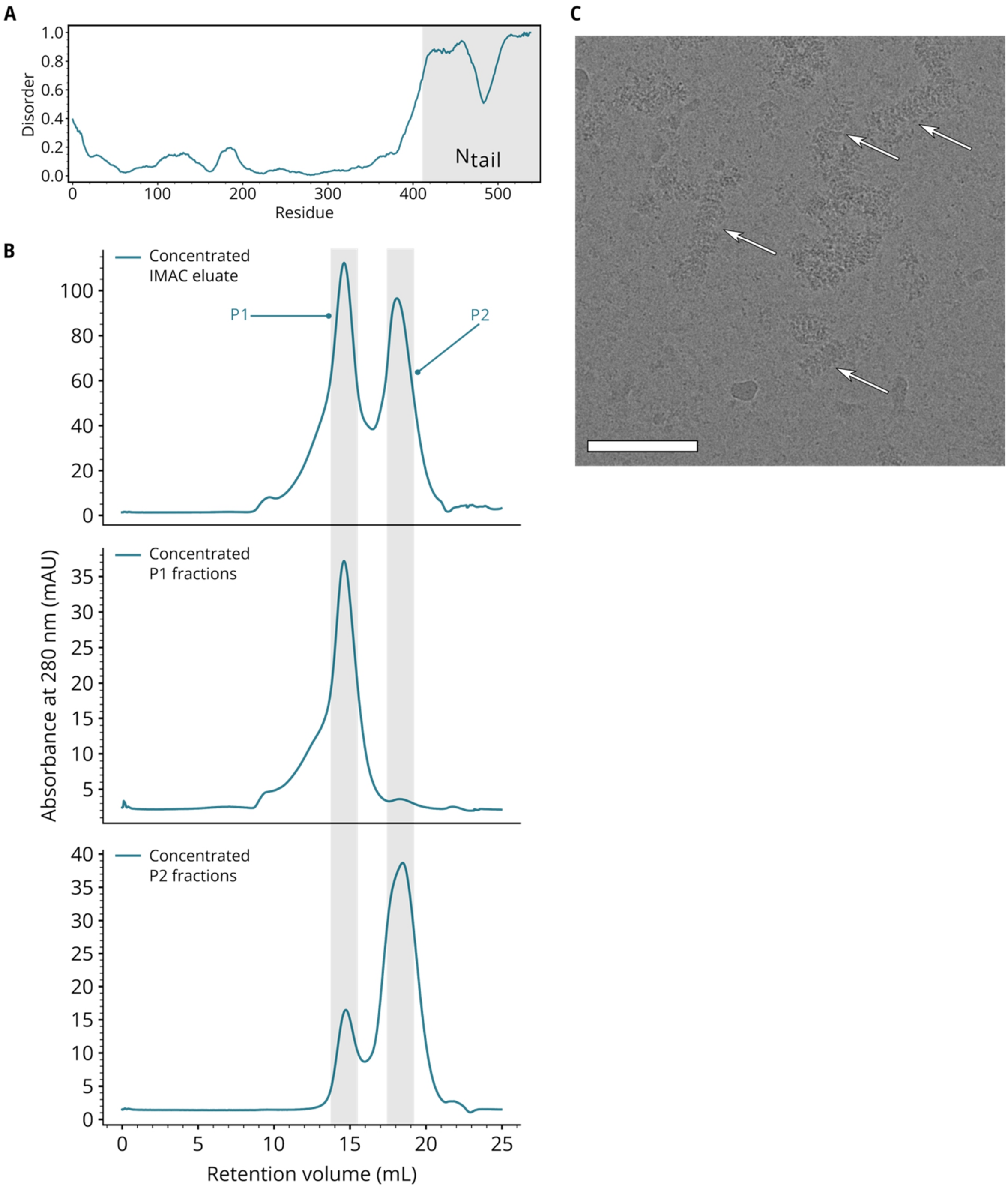


**Supplementary Figure 2: Purification of LayV-N and *in vitro* reconstitution of helical assemblies (A)** Disorder plot of LayV-N, predicted with Metapredict v3 (*1*), with the $N_{tail}$ residues highlighted. **(B)** Size exclusion chromatograms of LayV-$N_{core}$ purification. After immobilized metal affinity chromatography, the LayV-$N_{core}$ eluate was concentrated and subjected to size exclusion chromatography (SEC). From this run, two populations (P1, oligomer, and P2, monomer) of the protein were observed. Both these populations were concentrated separately and subjected to SEC individually (original purification: top, P1 fraction only: middle, P2 fraction only: bottom). **(C)** Representative cryo-EM micrograph of the *in vitro* reconstituted LayV-$N_{core}$ P2 + PolyA-$RNA_6$ sample. Helical assemblies are highlighted with white arrows. Scale bar = 100 nm.

Begränsad delning

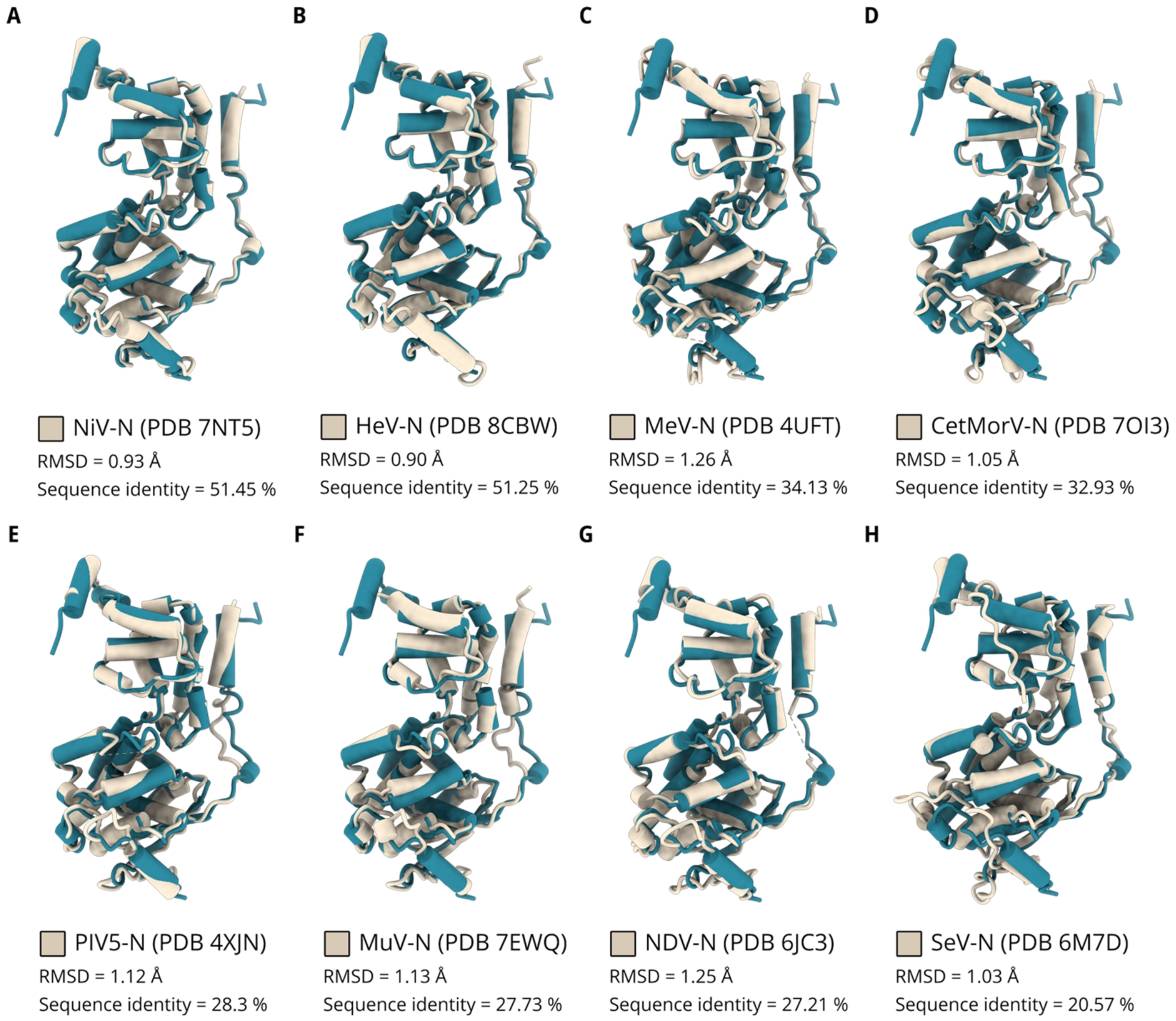


**Supplementary Figure 3: Structural alignment of LayV-N-closed with other available closed N structures of *Paramyxoviridae*.** Structure of LayV-N-closed from this study (coloured in teal, depicted here in cartoon representation (cylinders)) was aligned with the available paramyxoviral closed N structures (coloured in beige in respective panels) in UCSF ChimeraX 1.7.1 (*2*). LayV-N-closed was aligned with **(A)** NiV-N (PDB 7NT5) (*3*), **(B)** HeV-N (PDB 8CBW) (*4*), **(C)** MeV-N (PDB 4UFT) (*5*), **(D)** CetMorV-N (PDB 7OI3) (*6*), **(E)** PIV5-N (PDB 4XJN) (*7*), **(F)** MuV-N (PDB 7EWQ) (*8*), **(G)** NDV-N (PDB 6JC3) (*9*), and **(H)** SeV-N (PDB 6M7D) (*10*). All closed N structures aligned with LayV-N-closed are depicted as cartoons (cylinders; in beige). RNA omitted for clarity. (NiV = Nipah virus, HeV = Hendra virus, MeV = Measles virus, CetMorV = Cetacean morbillivirus, PIV5 = Parainfluenza virus 5, MuV = mumps virus, NDV = Newcastle disease virus, and SeV = Sendai virus).

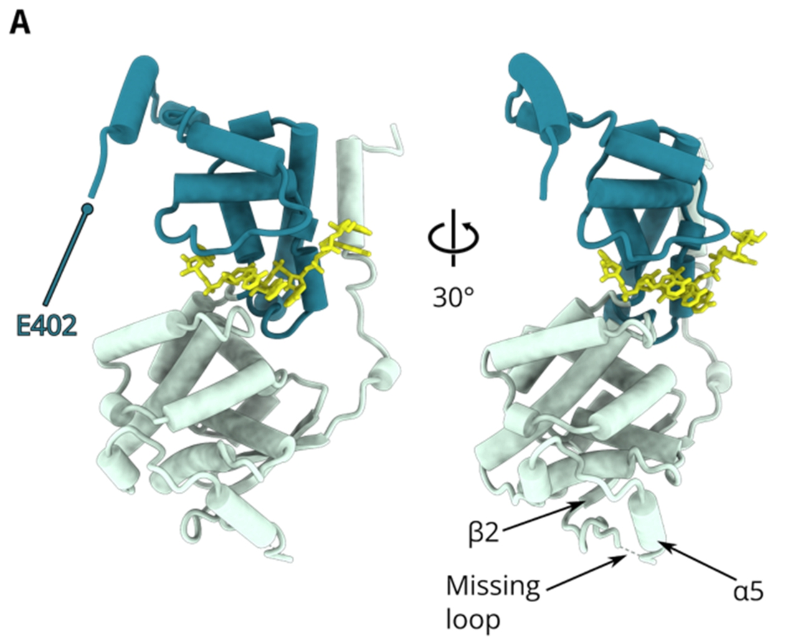


**Supplementary Figure 4: Unmodelled regions of LayV-N-closed. (A)** LayV-N-closed model shown here in a cartoon representation (cylinders) with the NTD and CTD domains coloured in green and teal, respectively. PolyA-$RNA_6$ is shown as sticks. A portion of the loop connecting beta sheet β2 and alpha helix α5 was disordered and thus unmodelled. Similarly, the density map for C-terminus was disordered from residue 403 onward.

Begränsad delning

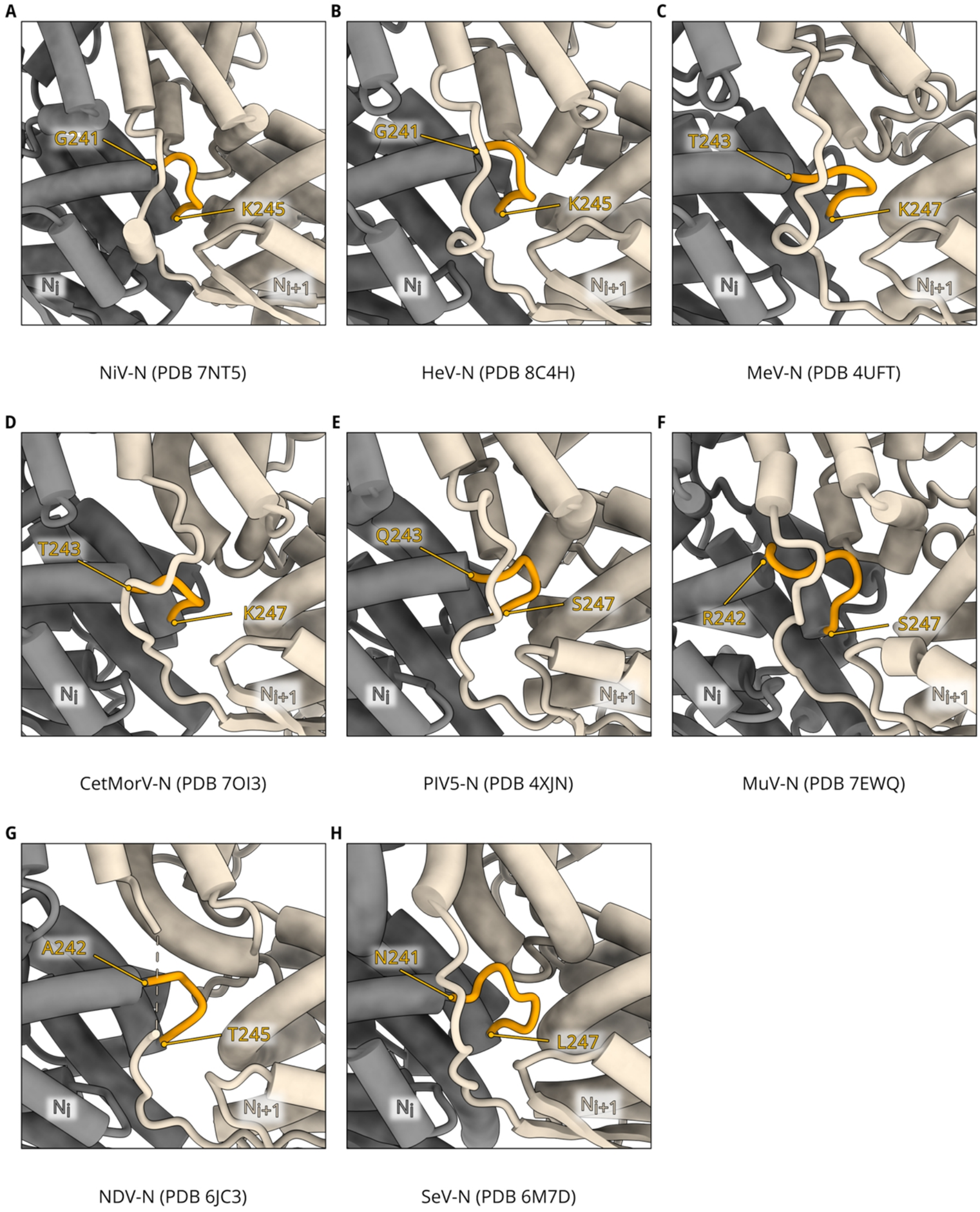


**Supplementary Figure 5: '$N_{NTD}$ loop – N-hole insertion' interaction observed in other paramyxoviruses.** Closeups of lateral interface between $N_i$ (grey) and $N_{i+1}$ (beige) protomers, focused on the insertion of NTD-loop (orange) into N-hole. Residue boundaries of the NTD loop are indicated. Structures of closed N shown here are from **(A)** NiV-N (PDB 7NT5) (*3*), **(B)** HeV-N (PDB 8CBW) (*4*), **(C)** MeV-N (PDB 4UFT) (*5*), **(D)** CetMorV-N (PDB 7OI3) (*6*), **(E)** PIV5-N (PDB 4XJN) (*7*), **(F)** MuV-N (PDB 7EWQ) (*8*), **(G)** NDV-N (PDB 6JC3) (*9*), and **(H)** SeV-N (PDB 6M7D) (*10*). All N structures are represented as cartoons (cylinders).

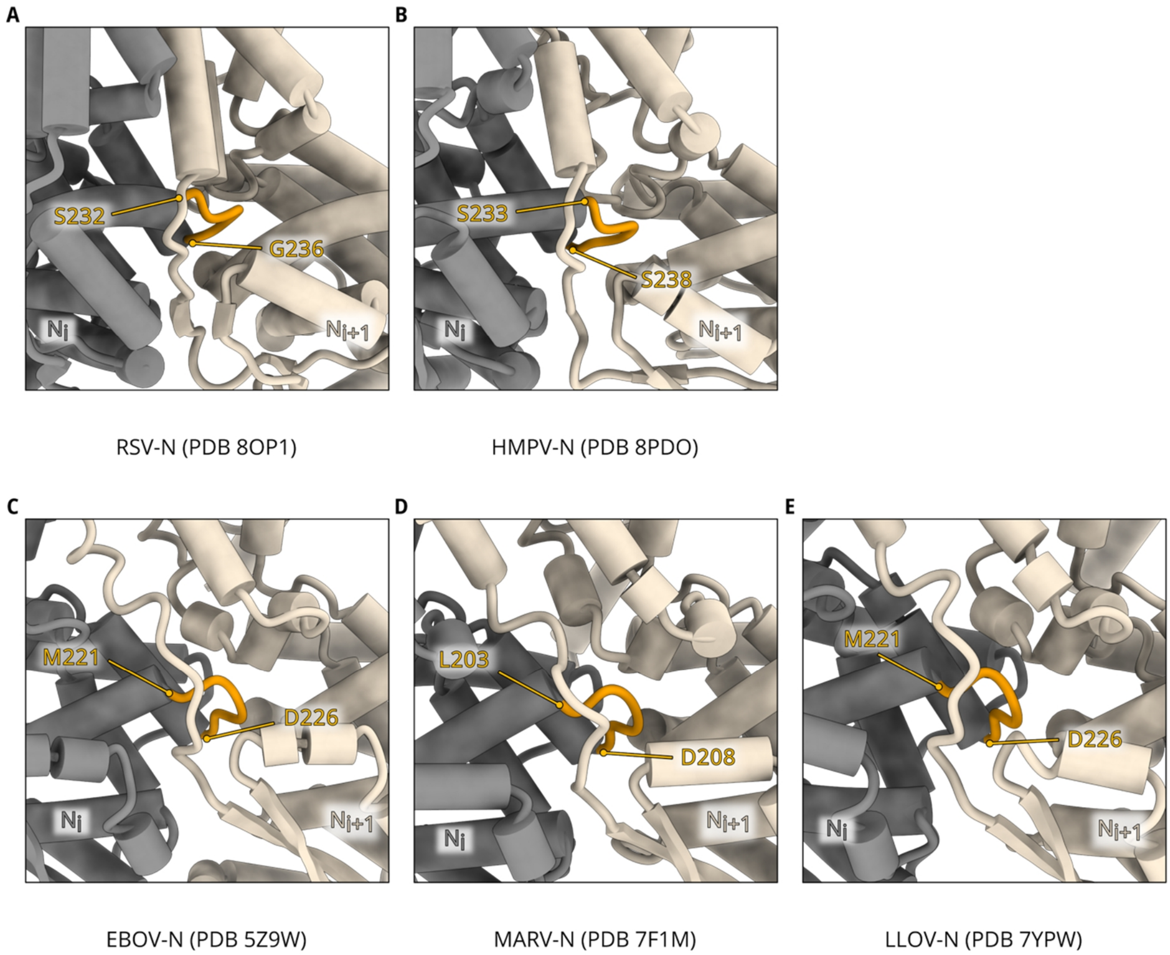


**Supplementary Figure 6: '$N_{NTD}$ loop – N-hole insertion' interaction observed in pneumoviruses and filoviruses.** Closeups of lateral interface between $N_i$ (grey) and $N_{i+1}$ (beige) protomers, focused on the insertion of NTD-loop (orange) into N-hole. Residue boundaries of the NTD loop are indicated. Structures of closed N shown here are from **(A)** RSV-N (PDB 8OP1) (*11*), **(B)** HMPV-N (PDB 8PDO) (*12*), **(C)** EBOV-N (PDB 5Z9W) (*13*), **(D)** MARV-N (PDB 7F1M) (*14*), **(E)** LLOV-N (PDB 7YPW) (*15*). All N structures are represented as cartoons (cylinders). (RSV = Respiratory syncytial virus, HMPV = Human metapneumovirus, EBOV = Ebola virus, MARV = Marburg virus, and LLOV = Lloviu cuevavirus).

Begränsad delning

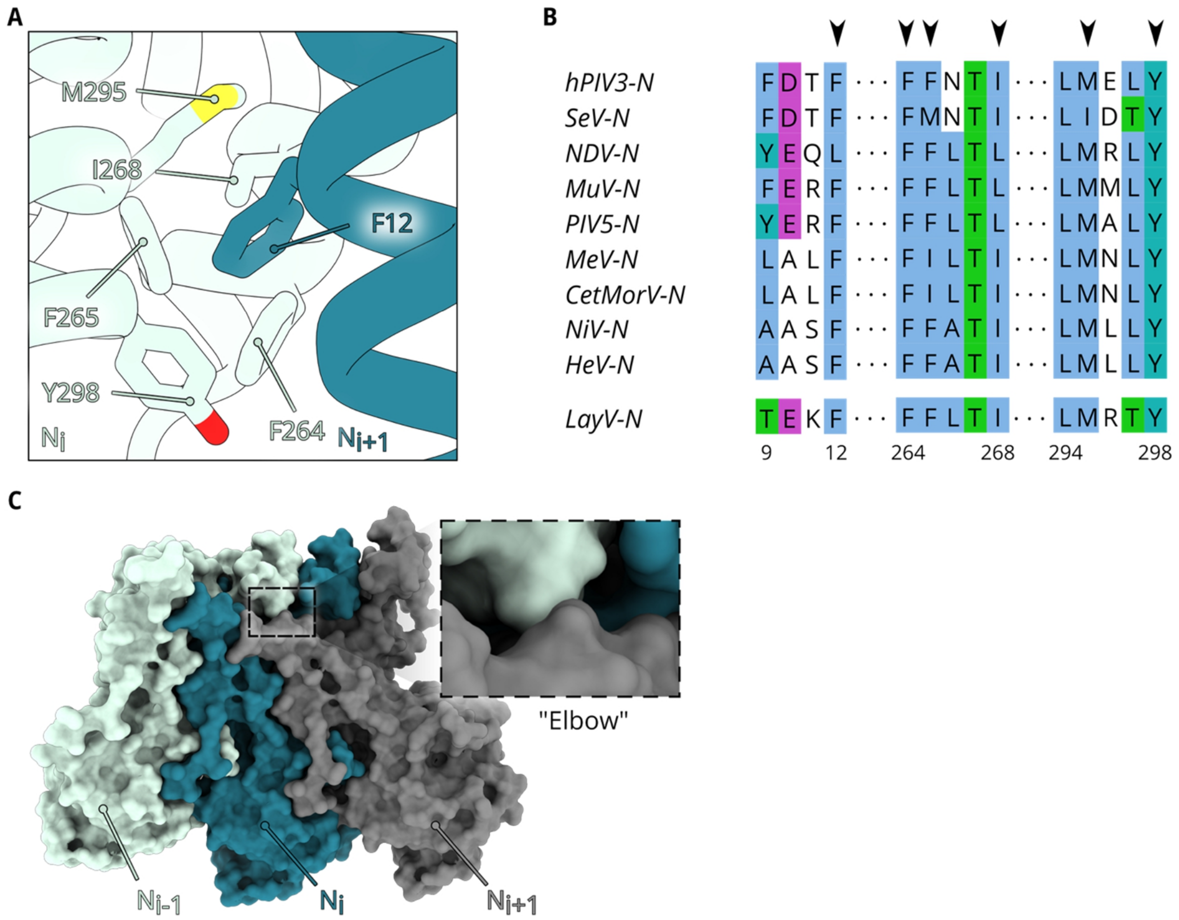


**Supplementary Figure 7: Lateral interface interactions of LayV-N$_{core}$ helical assembly. (A)** A cartoon representation of the interactions mediated by F12 residue in NT-arm of $N_{i+1}$ protomer with $N_{NTD}$ of $N_i$ protomer. Sidechains of residues are depicted as sticks. **(B)** Multiple sequence alignment of paramyxoviruses showing the conserved nature of residues involved in the interaction shown in (A): amino acids 12, 264, 265, 268, 295, and 298 (Black arrows; Residue numbering based on LayV-N). For clarity, only the regions of interest are shown. (hPIV3 = Human parainfluenza virus 3). **(C)** Surface representation of three consecutive protomers $N_{i-1}$, $N_i$, and $N_{i+1}$ (coloured in green, teal, and grey, respectively) from the helical assembly with a closeup of the "elbow" interaction between the CT-arm of $N_{i-1}$ and NT-arm of $N_{i+1}$.

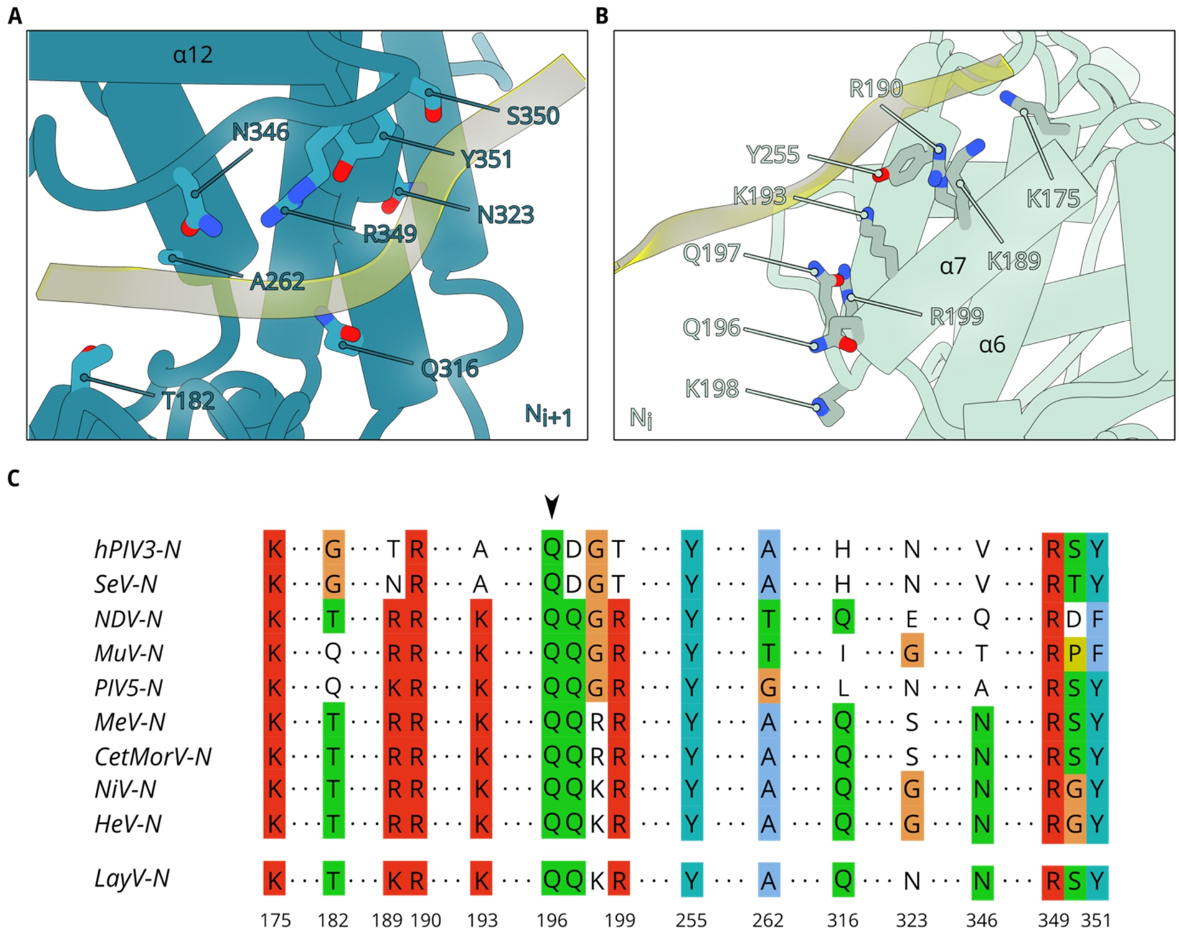


**Supplementary Figure 8: RNA binding residues of LayV-N.** Cartoon representations of the interactions mediated by PolyA-$RNA_6$ (shown as ribbon; coloured in yellow) with $N_{i+1}$ **(A)** and $N_i$ **(B)** protomers (depicted as cartoons; coloured in teal and green, respectively). Residues involved in the interaction are highlighted and shown as sticks. **(C)** Multiple sequence alignment of paramyxoviruses showing the conserved nature of residues involved in the interactions shown in (A) and (B): 175, 182, 189, 190, 193, 196, 199, 255, 262, 316, 323, 346, 349, and 351. Residue Q196 is highlighted with a black arrow. (Residue numbering based on LayV-N). For clarity, only the regions of interest are shown.

Begränsad delning

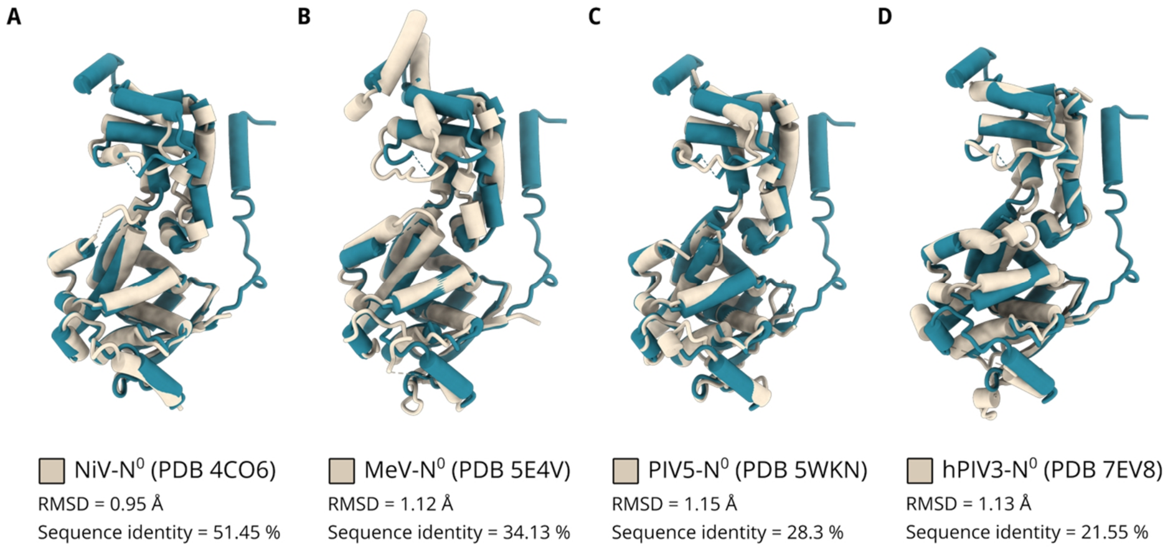


**Supplementary Figure 9: Structural alignment of LayV-N-opened with other available opened N (N$^{0}$-P) structures of *Paramyxoviridae*.** Structure of LayV-N-opened from this study (coloured in teal, depicted here in cartoon representation (cylinders)) was aligned with the available paramyxoviral N$^{0}$-P structures (coloured in beige in respective panels) in UCSF ChimeraX 1.7.1 (*2*). LayV-N-opened was aligned with N$^{0}$-P complexes of **(A)** NiV (PDB 4CO6) (*16*), **(B)** MeV (PDB 5E4V) (*17*), **(C)** PIV5 (PDB 5WKN) (*18*), and **(D)** hPIV3 (PDB 7EV8) (*19*). In all alignments shown here, N$^{0}$ structures aligned with LayV-N-opened are depicted as cartoons (cylinders) and the respective P residues have been omitted for clarity.

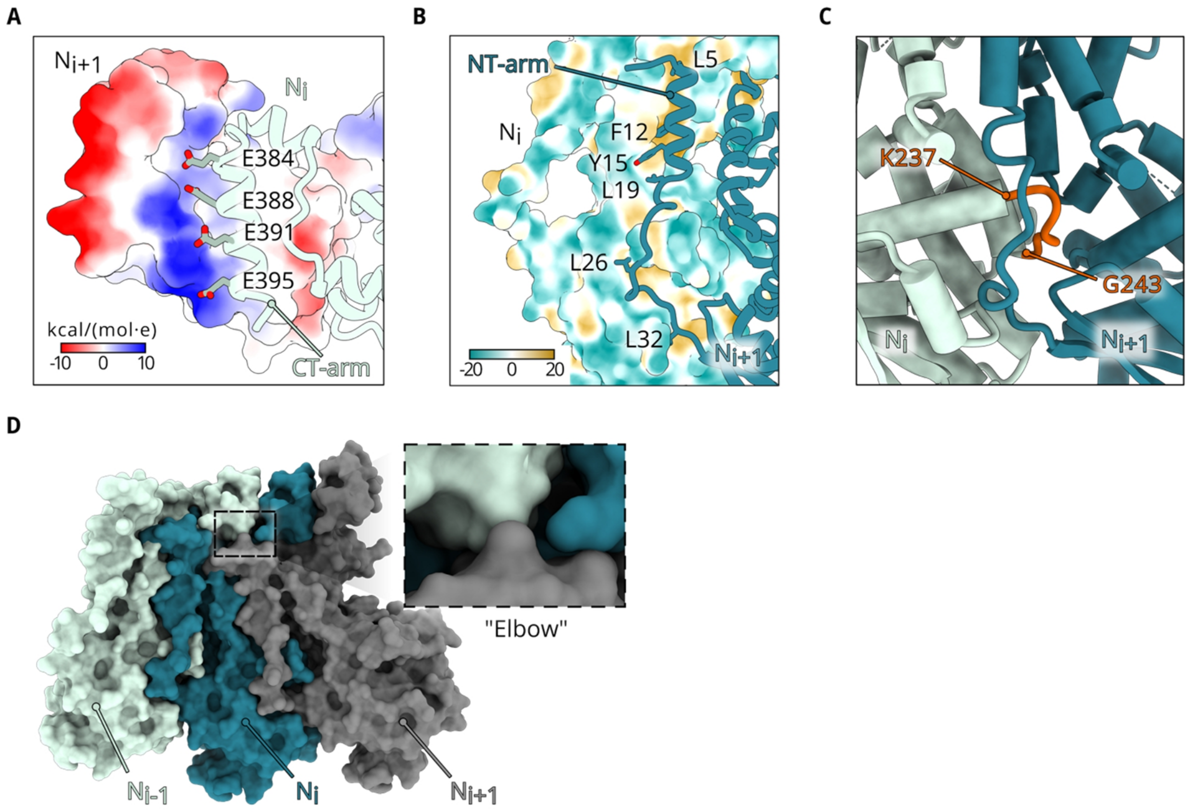


**Supplementary Figure 10: Lateral interface interactions of LayV-$N_{core}$ ring assembly. (A)** Closeup of the interface between CT-arm of $N_i$ (green ribbon) and the CTD of $N_{i+1}$ (surface representation, coloured by electrostatic potential as indicated). Glutamates of the CT-arm are shown as sticks. **(B)** Closeup of the interface between NT-arm of $N_{i+1}$ (teal ribbon) and the $N_i$ protomer, shown as surface and coloured by hydrophobicity (Brown = most hydrophobic, green = most hydrophilic). **(C)** Closeup of the lateral interface focused on the insertion of NTD-loop (orange) into N-hole. Residue boundaries of the NTD loop are indicated. **(D)** Surface representation of three consecutive protomers $N_{i-1}$, $N_i$, and $N_{i+1}$ (coloured in green, teal, and grey, respectively) from the helical assembly with a closeup of the “elbow” interaction between the CT-arm of $N_{i-1}$ and NT-arm of $N_{i+1}$.

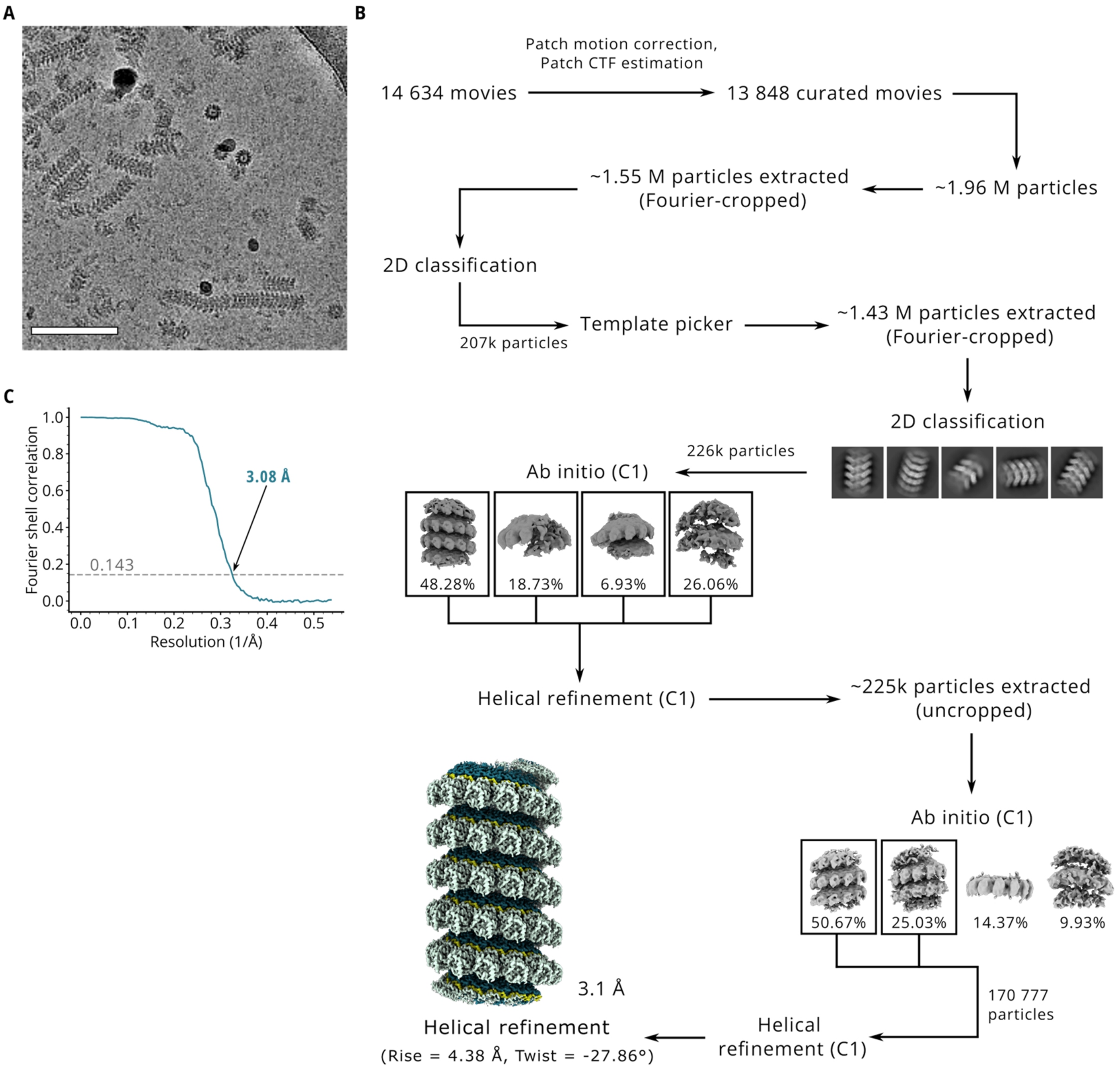


**Supplementary Figure 11: Cryo-EM image processing of LayV-N$_{core}$ helical assembly. (A)** A representative cryo-EM micrograph of the *in vitro* reconstituted LayV-N$_{core}$ P2 + PolyA-RNA$_6$ sample. Scalebar = 100 nm. **(B)** Cryo-EM image analysis pipeline followed for *in vitro* reconstituted helical assemblies of LayV-N$_{core}$ + PolyA-RNA$_6$. **(C)** Fourier shell correlation of the cryo-EM map of the helical assembly, plotted as a function of the spatial frequency. The map has a resolution of 3.1 Å based on the gold standard FSC of 0.143.

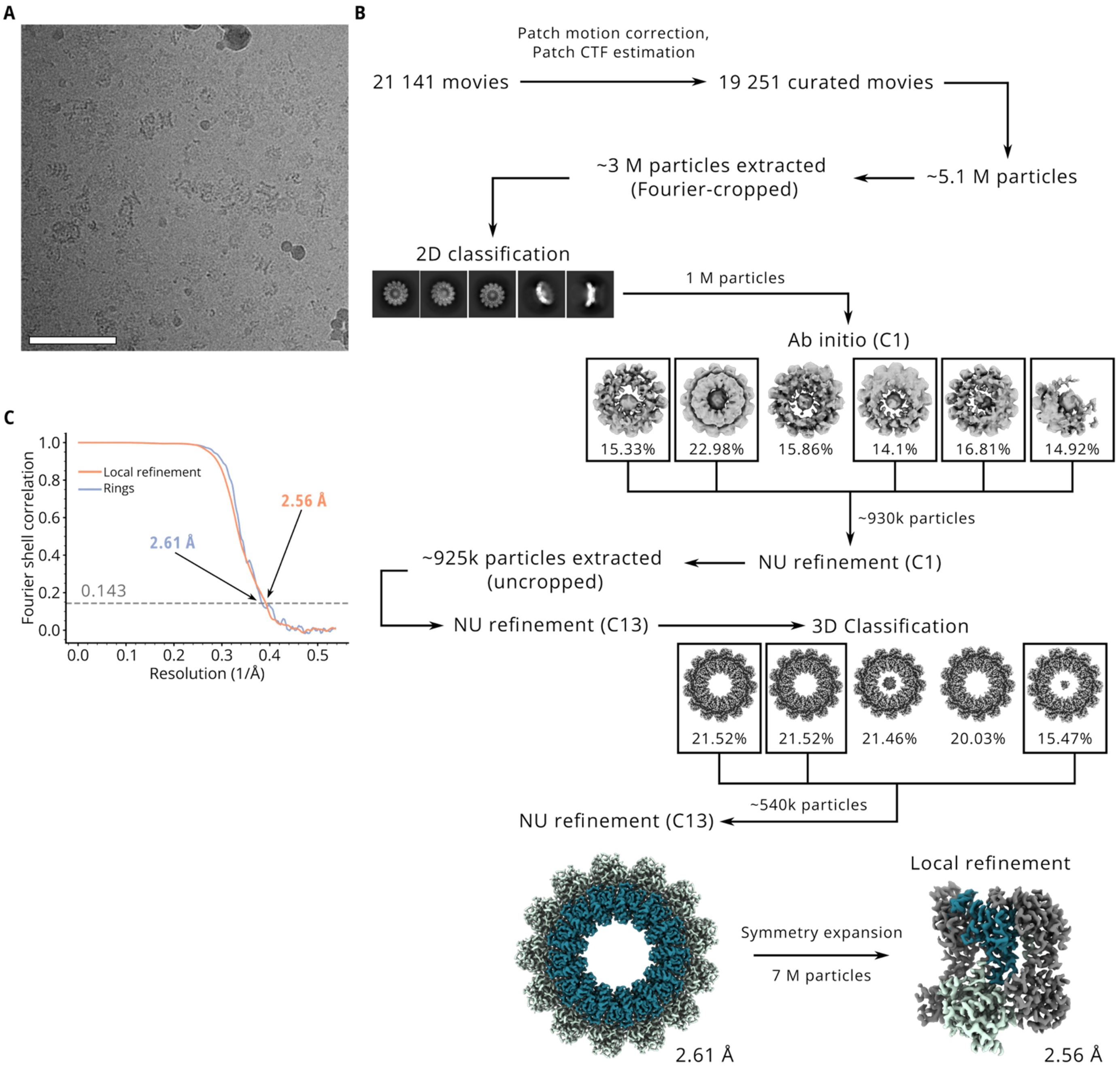


**Supplementary Figure 12: Cryo-EM image processing of RNA-free LayV-N$_{core}$ ring assembly. (A)** A representative cryo-EM micrograph of the LayV-N$_{core}$ P1 sample. Scalebar = 100 nm. **(B)** Cryo-EM image analysis pipeline followed for RNA-free ring assembly of LayV-N$_{core}$. **(C)** Fourier shell correlation for the cryo-EM maps of ring assembly and the local refinement, plotted as functions of the spatial frequency. The maps have resolutions of 2.61 Å and 2.56 Å, respectively, based on the gold standard FSC of 0.143.

Begränsad delning

Figure 3F:WT P2 + PolyA

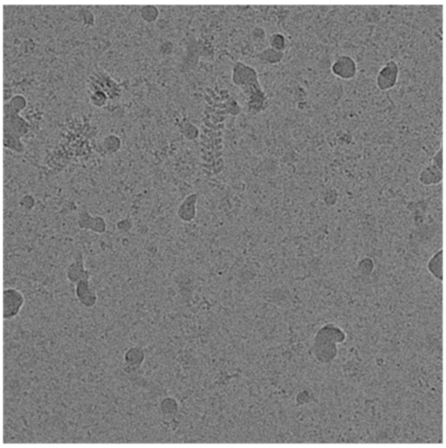

Figure 3F: R400E P2 + PolyA

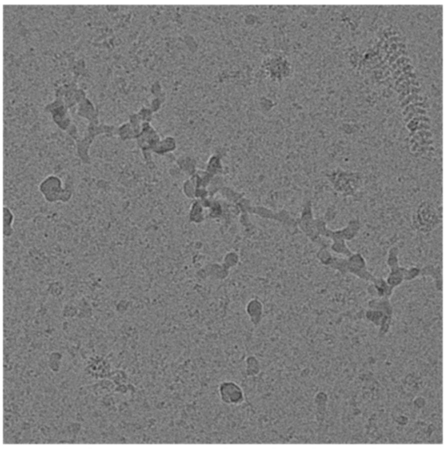

Figure 3F: E402F P2 + PolyA

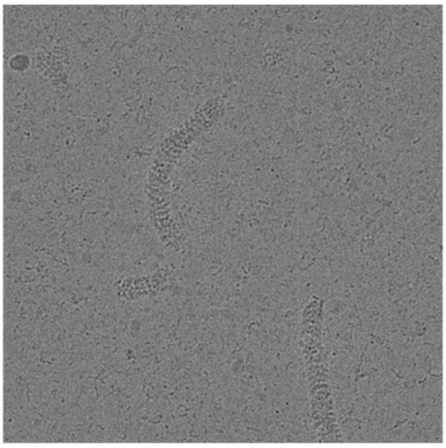

Figure 3F: E230R P2 + PolyA

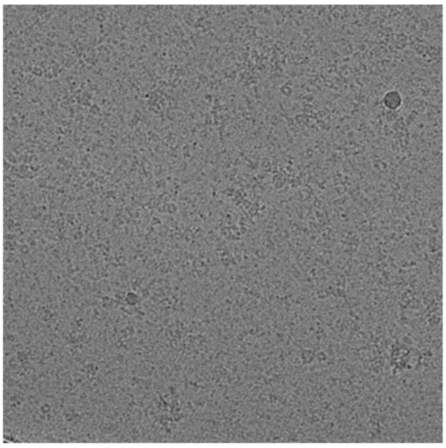

Figure 3F: E234F P2 + PolyA

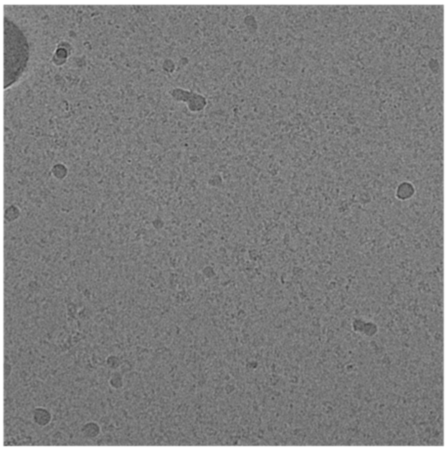

Figure 4C: LayV-N WT P1 + PolyA

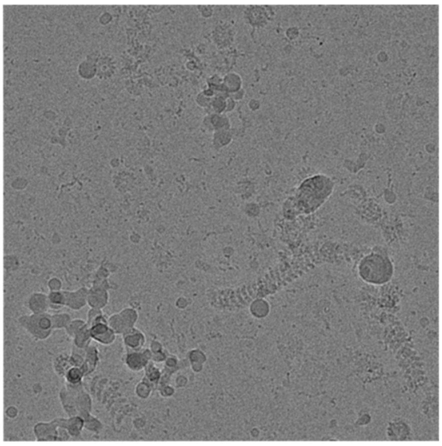

**Supplementary Figure 13: Uncropped Cryo-EM micrographs from figure panels 3F and 4C.**

**Supplementary Table 1: Lateral interface area between two consecutive protomers**

| | Lateral interface area (Å$^2$) |
|---|---|
| LayV-N (This study) | 3810 |
| SeV-N (PDB 6M7D) | 3300 |
| NiV-N (PDB 7NT5) | 3000 |
| HeV-N (PDB 8CBW) | 2930 |
| PIV5-N (PDB 4XJN) | 2900 |
| MeV-N (PDB 4UFT) | 2700 |
| NDV-N (PDB 6JC3) | 2700 |
| CetMorV-N (PDB 7OI3) | 2300 |
| MuV-N (PDB 7EWQ) | 2300 |

Begränsad delning

**Supplementary Table 2: Cryo-EM data acquisition, processing, and validation parameters**

| | LayV-N$_{core}$ NC-like complex<br>EMDB-XXXXX<br>PDB XXX | LayV-N$_{core}$ 13mer Ring<br>EMDB-XXXXX | LayV-N$_{core}$ Local refinement<br>EMDB-XXXXX<br>PDB XXX |
|---|---|---|---|
| **Data collection** | | | |
| Voltage (keV) | 300 | 300 | |
| Magnification | 130 000x | 130 000x | |
| Exposure rate ($e^-$/px/s) | 7.36 | 8.92 and 9.93 | |
| Total dose ($e^-$/Å$^2$) | 40 | 40 | |
| Nominal defocus range (μm) | -1.2 to -2.4 | -1.2 to -2.4 | |
| Pixel size (Å) | 0.92 | 0.92 | |
| Movies collected | 14 598 | 21 141 | |
| **Processing** | | | |
| Movies used | 13 848 | 19 251 | |
| Initial no. of particles | 1 546 278 | 2 991 660 | |
| Final no. of particles | 170 777 | 541 364 | 7 037 732 (sym. expansion) |
| Symmetry imposed | Helical<br>Rise = 4.38 Å<br>Twist = -27.86° | C13 | C1 |
| Final map resolution (Å) | 3.08 | 2.61 | 2.56 |
| FSC threshold | 0.143 | 0.143 | 0.143 |
| *B* factor (Å$^2$) | -55.7 | -122.1 | -86.3 |
| **Refinement** | | | |
| Model composition | | | |
| Amino acids | 398 | | 381 |
| Nucleotides | 6 | | 0 |
| Model resolution (Å) | 3.2 | | 2.9 |
| (Model-Map FSC) | 0.5 | | 0.5 |
| ADP (Å$^2$) | | | |
| Protein | 102.91 | | 77.45 |
| RNA | 69.74 | | - |
| R.m.s deviations | | | |
| Bond lengths (Å) | 0.001 | | 0.002 |
| Bond angles (°) | 0.394 | | 0.402 |
| **Validation** | | | |
| Molprobity score | 1.37 | | 1.30 |
| Clashscore | 6.71 | | 5.57 |
| Rotamer outliers (%) | 0.29 | | 0.61 |
| Model vs map CC | 0.85 | | 0.82 |
| Ramachandran plot | | | |
| Favoured (%) | 98.98 | | 98.66 |
| Allowed (%) | 1.02 | | 1.34 |
| Outliers (%) | 0.00 | | 0.00 |
